# Comprehensive analysis of non-synonymous single-nucleotide polymorphism of human TSC1 and TSC2 genes: an in silico approach

**DOI:** 10.64898/2026.02.04.703811

**Authors:** Tasnim Alam, Shangida Akther

## Abstract

Tuberous sclerosis complex (TSC) is an autosomal dominant disorder caused by mutations in the TSC1 and TSC2 genes and is characterized by benign hamartoma formation in multiple organs. The TSC1–TSC2 complex regulates mTORC1 signaling in response to cellular growth conditions. This study aims to predict the structural stability and functional effects of non-synonymous single-nucleotide polymorphisms (nsSNPs) in human TSC1 and TSC2 using computational approaches. Twelve computational tools were assessed using receiver operating characteristic (ROC) analysis and applied to identify deleterious nsSNPs. Protein stability was predicted using I-Mutant 2.0 and MUpro, while evolutionary conservation was analyzed with ConSurf. NetPhos 3.1 identified potential PTM sites, and MutPred2.0 evaluated their functional impact. Project HOPE assessed mutation-induced physicochemical changes. Structural models were validated using multiple tools, visualized in ChimeraX 1.9, and further evaluated by molecular dynamics simulation to confirm wild-type and mutant stability. All twelve tools had AUC values above 0.90. A combined in silico analysis identified twelve high-risk nsSNPs in TSC1 and sixteen in TSC2, all reducing protein stability, located in conserved regions, and potentially disrupting phosphorylation sites. MutPred and Project HOPE confirmed their impact on protein function. Functional analysis showed TSC1 and TSC2 affect mTORC1 and PI3K–Akt pathways. RMSF and RMSD analyses revealed that TSC1 variants rs1846545280 (G236E), and rs2132135678 (V234E), and TSC2 variants rs45517223 (S758C), rs2151354925 (T836P), and rs45517365 (R1570W) had the largest structural fluctuations. Substitution with glutamic acid, a negatively charged and bulkier residue, may disrupt local folding of TSC1. Similarly, replacement of arginine with tyrosine at position 1570 may impair Rheb binding at the GAP domain of TSC2. These findings highlight potentially pathogenic nsSNPs in TSC1 and TSC2.

## Introduction

Tuberous Sclerosis Complex (TSC) is a multisystemic disorder and autosomal dominant tumor syndrome characterized by the development of numerous benign tumors (e.g. hamartomas) most commonly affecting the brain, kidneys, skin, heart, and lungs (1). Functionally, the TSC1/TSC2 complex is positioned at the center of multiple growth signalling pathways and is a key integrator of signals controlling protein translation and cell growth (2). The TSC complex consists of the hamartin (encoded by TSC1), the tuberin (TSC2 gene product), and the recently identified TBC1D7, which forms high oligomeric assemblies in cells (3). Structurally, hamartin (TSC1) and tuberin (TSC2) exhibit distinct functional domains that underpin their regulatory roles in cell growth and mTORC1 signaling. TSC1 contains an N-terminal α-helical/HEAT repeat region, a putative transmembrane segment, and extensive coiled-coil regions in its central and C-terminal portions that mediate heterodimerization with TSC2 and complex assembly with TBC1D7 (4). Additionally, N-terminal region of TSC1 has been displayed to do Rho GTPase regulation, contributing to cytoskeletal dynamics and focal adhesion(5). TSC2 is characterized by an extended HEAT repeat/α-solenoid domain at the N-terminus that interacts with TSC1, followed by a dimerization domain (DD), and a GTPase-activating protein (GAP) domain (6). Although the GAP activity of TSC2 is of obvious functional importance to the complex, TSC1 is required to stabilize TSC2 and prevent its ubiquitin-mediated degradation (7). TSC2 has also been shown to physically interact with the transcription factor FoxO1 via a C-terminal region linking TSC2 to insulin signaling pathways and functional modulation of the TSC1-TSC2 complex (8). Aside from those affecting the TSC2 GAP domain, missense mutations in TSC1 or TSC2 commonly destabilize the complex, thereby leading to TSC2 degradation (9). There are different forms of SNPs, and a missense SNP, which is a type of non-synonymous SNP (nsSNP) substitution characterized by amino acid substitution with the possibility of forming a mutated protein with new structural and functional features, is the most important form of SNPs (10). This form of SNP can produce deleterious actions by minimizing protein solubility, destabilizing protein tertiary structure, and manipulating gene regulation through modifying transcription regulatory proteins, furthermore causing cancer (11,12)Single-nucleotide polymorphisms (SNPs) account for more than 90% of all nucleic acid sequence variations in humans (13). While a large number of missense SNPs for several genes have been reported and deposited in databases, the total number of missense SNPs for each gene showed a variation in the attribution to disease modification; therefore, it became essential to filter the SNPs with possible pathogenicity from the pool of neutral variants (14).

Comprehensive screening for mutations at both the TSC1 and TSC2 loci has been performed in several large cohorts of TSC patients, and a wide variety of different pathogenic mutations have been discovered. Although ∼20% of the mutations identified in the TSC2 gene are missense changes, missense mutations in the TSC1 gene appear much less frequently. To characterize pathogenic variants, mutational and molecular analysis had been performed in several different demographics such as Mexican (15), Brazilian (16), Chinese (17), Danish (18), Japanese (19), and Greek (20). The missense variants resulted in significantly reduced TSC1 and TSC2 signals and also were unable to inactivate TORC1-dependent S6K-T389 phosphorylation, similar to the pathogenic p.R611Q variant (21). Three TSC2 mutants (R505Q, H597R, and L1624P) showed reduced affinity in binding to TSC1, perhaps due to conformational changes or because of increased targeting of TSC2 for degradation (22). Also missense variants of TSC2 were found to cause a significant reduction in the ability of the TSC complex to inhibit S6K-T389 phosphorylation. The p.S1045F and p.I1197F substitutions affect the TSC complex, thus increasing mTOR activity (23). Several TSC1 variants have been associated with different cancers. TSC1 rs7874234 is hypothesized to be functional in ER+ breast cancer because the T allele, but not the C allele, may create an estrogen receptor element (ERE) site, resulting in increased TSC1 transcription and subsequent inhibition of mTOR (24). Missense variants present in the N-terminal domain of TSC1, were found to be associated with bladder cancer (2). The TSC1 rs13295634 (TG/GG) allele has been associated with the worst survival in colorectal cancer (25). A missense variant p.Ser208Gly of TSC1 is found to have a damaging impact in keratoconus patients (26).

The TSC1 gene is less studied than TSC2 as most of the mutations causing the disease are found in TSC2 though TSC1 is equally important in the formation of TSC complex. In this context, computational approaches provide a cost-effective and time-efficient alternative to preliminary experimental screening, enabling the prioritization of functionally significant SNPs that may disrupt TSC complex stability or Rheb binding for subsequent experimental validation. This study aims to characterize functionally significant SNPs using in silico approaches, serving as an initial step toward identifying the most deleterious variants of both TSC1 and TSC2 in the human genome.

## Methodology

nsSNPs data was retrieved from the NCBI-dbSNP database. To predict the most deleterious SNP, multiple computational tools of varying principles were employed. Finally, the predicted SNPs were analyzed for their role in protein stability.

### 2.1 Retrieving nsSNP data of TSC1 and TSC2 gene

The National Center for Biotechnology Information, dbSNP database, was used to retrieve the nsSNPs of the TSC1 and TSC2 genes (https://www.ncbi.nlm.nih.gov/snp/). The sequences of both protein Q92574 and P49815 were obtained from the UniProtKB database (https://www.uniprot.org/). During dataset preparation, only missense variants of nsSNPs were chosen because their impact can vary from benign to highly detrimental, depending on the role of the substituted amino acid in the protein’s function or structure compared to a nonsense variant. As a result, the coding regions of 2583 (TSC1) and 4468 (TSC2) nsSNPs were prioritized for further investigation due to their recognized ability to significantly influence protein structure and function. Subsequently, all the nsSNPs were isolated and retrieved for further investigation.

### 2.2 Accuracy of nsSNP prediction tools

Data from pathogenic and benign variants of TSC1 and TSC2 were used as experimental data to assess the accuracy of the nsSNP Prediction tools. Receiver operating characteristic (ROC) curves were generated using the pROC package in R (27). The area under the curve (AUC) was calculated to assess the predictive performance of each tool.

### 2.3 Predicting Deleterious Variants

As per the ACMG standards and guidelines, it recommends that multiple prediction software programs should be used for sequence variant interpretation as each in silico tools have their own weaknesses and strengths. A total of twelve tools were used to predict the damaging and deleterious nsSNPs of the both TSC1 and TSC2 genes: PROVEAN (Protein Variation Effect Analyzer), SIFT (Sorting Intolerant From Tolerant), PolyPhen-2 (Polymorphism Phenotyping v2), FATHMM (Functional Analysis Through Hidden Markov Models), AlphaMissense, Bayesdel (Bayesian Deleteriousness), MetaLR (Meta-Logistic Regression), CADD, REVEL, VEST4, ClinPred, M-CAP. The cut-off score of PROVEAN is 2.5 (27), while SIFT is ≤0.05 (28). A score above those numbers was considered benign. For PolyPhen2, the score varies between 0 and 1, where 0.45 to 0.95 is considered possibly damaging and 0.95 to 1.0 probably damaging (29). Variants with a FATHMM score below-1.5 are classified as deleterious (30). A score above a certain threshold (e.g., 0.56) suggests the variant is likely pathogenic, while a score below suggests it is likely benign in AlphaMissense (31). The cut-off score ranges from −1.29334 to 0.75731 for Bayesdel, and the higher the score is, the more deleterious the variant (32). The decision boundary to classify pathogenic or benign is 0.5 for ClinPred (lower for benign, higher for pathogenic) (33). MetaLR considers nsSNP as benign with scores of 0.5 or lower and vice versa (30). CADD has “raw score” and a scaled PHRED-like score that ranks the variant relative to all possible substitutions in the human genome. A higher score (>20) indicates a higher chance of deleteriousness (34). The cutoff score for REVEL is >0.5 (35). The VEST final score ranges from 0 to 1, and the higher the score is, the more deleterious the variant is (30). For M-cap, the higher the score is (>0.025), the more deleterious the variant is (36). nsSNPs predicted as “Disease/Deleterious” by all the twelve tools were picked for further analysis.

### 2.4 Stability prediction of nsSNPs on protein

I-Mutant2.0 and MUpro web server were utilized to estimate the potential effects of nsSNPs on the structural reliability of the protein and free energy change Delta Delta G (DDG). Accordingly, stability will increase when DDG is > 0 kcal/mol and decrease when DDG is < 0 kcal/mol. In this project, protein sequence interface of I-Mutant2.0 was used. Protein sequence, AA position, Residue change was given as input. Output is displayed as either free energy change or reliability index at pH 7.0 and temperature 25^0^ indicating ideal environment with the statement increasing or decreasing stability of protein (37). Both Support Vector Machines and Neural Networks are used in MUpro. After analysis, MUpro provides a confidence score ranging from-1 to +1, where a negative score indicates a predicted decrease in protein stability and a positive score suggests increased stability. Results from both SVM and Neural networks were incorporated into the project(38). Both types of results in the case of decreasing stability were integrated in this study.

### 2.5 Evolutionary Conservation Analysis of the most Damaging nsSNPs

Conservation surface map (ConSurf) is a computational tool designed for mapping evolutionarily conserved residues onto the surface of protein structures, providing insight into functionally or structurally important sites. It utilizes multiple sequence alignments (MSA). Residues essential for maintaining the protein’s fold and function tend to be highly conserved, while variable regions tolerate substitutions more freely. The final output is a color-coded visualization of the protein structure, typically ranging from highly conserved (colored dark purple or blue) to highly variable (colored cyan or turquoise). These mapped results help identify critical residues involved in structural stability, active sites, ligand binding, or protein-protein interaction interfaces, offering valuable guidance for functional annotation and mutational analysis (39).

### 2.6 Prediction of Post-Translational Modification (PTM) Sites

NetPhos 3.1 is a computational tool used to predict potential phosphorylation sites on protein sequences, identifying serine (S), threonine (T), and tyrosine (Y) residues that may undergo phosphorylation employing an artif icial neural network (ANN)–based algorithm. The tool assigns a phosphorylation probability score (0 to 1) to each site, with a higher score (>0.5) indicating a greater likelihood of modification (40).

### 2.7 Identification of Structural and Functional Alterations

MutPred2 uses a neural network–based algorithm trained on a large dataset of 53,180 pathogenic variants and 206,946 neutral variants from HGMD, SwissVar, and dbSNP. The tool outputs a g score (ranging from 0 to 1), where values >0.5 indicate a higher probability of the variant being deleterious. Additionally, it provides empirical p-values for various potential molecular consequences (e.g., loss of phosphorylation, altered secondary structure), indicating whether the mutation is likely to cause significant changes in specific biological properties (41).

### 2.8 Identifying Single Nucleotide Polymorphism Impact on Protein Structure

HOPE server was used to analyze the SNPs’ impact on the protein 3D structure. The HOPE server depends on several sources for collecting the needed information in addition to building homology models using the YASARA program to perform the required function (42). It investigates how changes in amino acid composition affect native structures and the variations in hydrophobicity, charge, and size between residues of the wild-type and mutant-types. Uniprot provides the TSC1 and TSC2 protein sequences to Project HOPE for predicting the 3D structure of the mutated protein.

### 2.9 Functional Enrichment Analysis

Functional enrichment analysis was performed using the enrichR package to detect statistically noteworthy (P < 0.05) associations between the input gene set and curated databases of disease (DisGenet), and complementary functional pathway (KEGG) (43). RStudio (4.2.2) was employed for this analysis. The GGplot2 library of R was used for visualization.

### 2.10 Prediction of Protein-Protein Interaction Network

The online database STRING (Search Tool for Recurring Instances of Neighboring Genes) predicts the PPI network of the TSC1 and TSC2 protein (https://string-db.org/). It retrieves the genes that are tangentially (via other genes) linked to the query gene using an evolutionary method. The STRING output furnishes details regarding the protein’s expression, localization, transcription, experimental evidence, and graphical depictions of the interactions in which the queried protein participates (44). Network was visualized by Cytoscape software.

### 2.11 Comparative 3D Modeling of TSC1, TSC2 and Its Mutant

Alphafold, Phyre2, and Swiss-Model were used to generate wild-type protein structures of TSC1 and TSC2. The Alphafold TSC1 structure was used as the wild type (45). The fold recognition method was adapted to model the 3D structure of the Tuberin domain of TSC2 due to the absence of a homologous structure. Phyre2 was used to generate the three-dimensional structure of the Tuberin domain (46). The template, c4jZ7A, was used by Phyre2 for the estimation of three-dimensional models and then, the visualization of the structures was done by ChimeraX 1.6. Swiss-Model modeled the Gap domain as close homologs were available (47). All models were evaluated with ERRAT, MolProbity, Verify 3D, and PROCHECK (48). Mutant protein structures were generated using ChimeraX 1.6, a molecular visualization and modeling tool that allows interactive mutagenesis (49). After substituting the wild-type amino acid with the desired mutant residue, the local conformation of the mutant side chain was optimized using the Dunbrack 2010 rotamer library. This step samples the most favorable side-chain conformations (rotamers) for the new residue based on statistically preferred dihedral angles from high-resolution crystal structures, minimizing steric clashes and unfavorable torsions. Once the mutant model was built and side-chain conformation optimized, mutant protein structure was superimposed onto the native (wild-type) protein structure using ChimeraX’s structural alignment tool. This superimposition allowed calculation of the Root Mean Square Deviation (RMSD) — a quantitative measure of the structural similarity between the two structures. A lower <0.2 value indicates stable, realistic mutant model. Each of these models has been energy-minimized by GalaxyWEB (https://galaxy.seoklab.org/) to relax the structure and remove steric constraints and verified for stereochemical quality (50).

### 2.12 Molecular Dynamics Simulation

GROMACS is a high-performance molecular dynamics (MD) simulation software used to model the physical movements of atoms and molecules over time, based on classical Newtonian mechanics (51). Initially, each protein was placed within a simulation box solvated with explicit water molecules (TIP3P water models) and neutralized with appropriate counter-ions (Na⁺ and Cl⁻) to replicate physiological conditions. The system underwent energy minimization to eliminate unfavorable steric clashes and optimize atomic positions by achieving a local minimum in the potential energy landscape. Following this, two equilibration phases were conducted: NVT equilibration, maintaining a constant number of particles, volume, and temperature to stabilize the thermal environment, and NPT equilibration, keeping the number of particles, pressure, and temperature constant to stabilize system pressure and density while allowing solvent and ions to adjust around the protein structure. Finally, a 100-nanosecond production MD run was carried out, during which atomic forces were calculated and Newton’s equations of motion integrated at each time step (typically 1–2 femtoseconds). Atomic positions, velocities, and energies were recorded at defined intervals, generating trajectory files for subsequent analysis of structural deviations, flexibility, and stability. Root Mean Square Deviation (RMSD) analysis was then used to monitor structural deviations from the initial conformation, with stable RMSD profiles indicating structural integrity. Further Root Mean Square Fluctuation (RMSF) analysis was used to directly compare the dynamic flexibility of native and mutant proteins under simulated physiological conditions.

## Result

### 3.1 Retrieving missense variants of nsSNPs of TSC1 and TSC2 genes

The NCBI dbSNP database was employed to get polymorphism data for the TSC1 and TSC2 genes. There were 25,875 SNPs in TSC1, out of which 1,091 are synonymous variants, 19,690 intron region variants, 2583 missense variants of nsSNPs, and the remaining were different types. There were 33,258 SNPs in TSC2, containing 2,048 synonymous variants, 25,743 intron region variants, 4468 missense variants of nsSNPs, and the remaining were different types (Fig 1). All other SNPs were excluded except 2583 (TSC1) and 4468 (TSC2) non-synonymous SNPs (missense variations) for this project. A missense variant is a single nucleotide change in the coding region for a different amino acid substitution, potentially affecting the protein’s structure and function.

**Fig 1:**
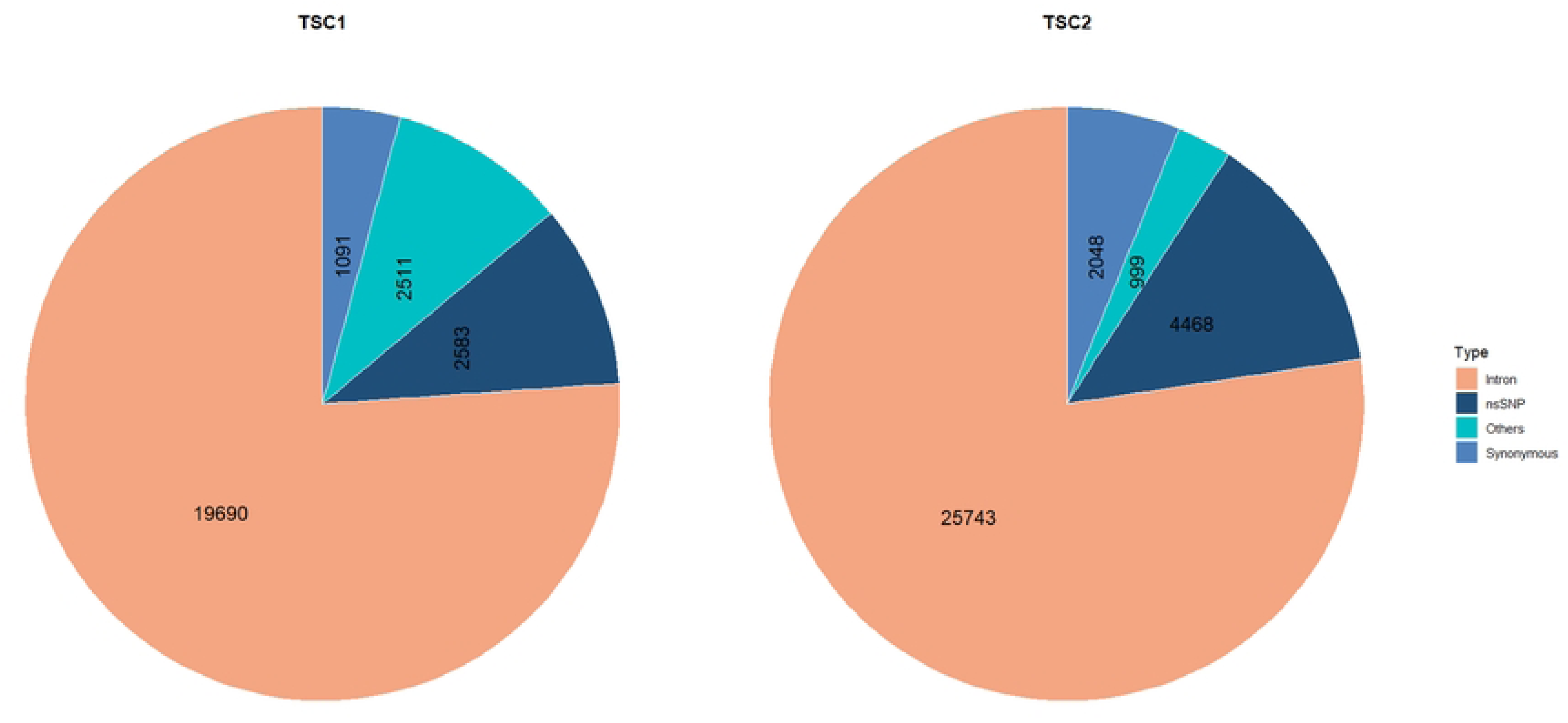
Piechart displaying each type of SNP in both TSC1 and TSC2. In TSC1, there are 2583 non-synonymous SNP, 1091 synonymous SNP, 19,690 intron and 2511 other. 4468 non-synonymous SNP, 2048 synonymous, 25473 intron, and 999 others in *TSC2*.

### 3.2 Assessing the specificity of in-silico pathogenicity prediction tools

The performance of twelve computational tools in discriminating between pathogenic and benign variants was evaluated using the Receiver Operating curve (ROC). The overall identification ability of each tool was quantified by the Area Under the Curve (AUC). The tools demonstrated high performance, with all AUC values exceeding 0.90. Tools with the highest AUC scores were ClinPred (AUC: 0.979), mcap (AUC: 0.977), and Revel (AUC: 0.976). These three tools exhibit highest specificity to rank pathogenic variants than benign variants compared to the other predictors such as Bayesdel (AUC: 0.975), AlphaMissense (AUC: 0.974), Sift (AUC: 0.915), CADD (AUC: 0.909), Fathmm (AUC: 0.937), Polyphen-2 (AUC: 0.948), Vest4 (AUC: 0.967), MetaLR (AUC: 0.951) and lastly Provean (AUC: 0.974) (**Fig 2**).

**Fig 2:**
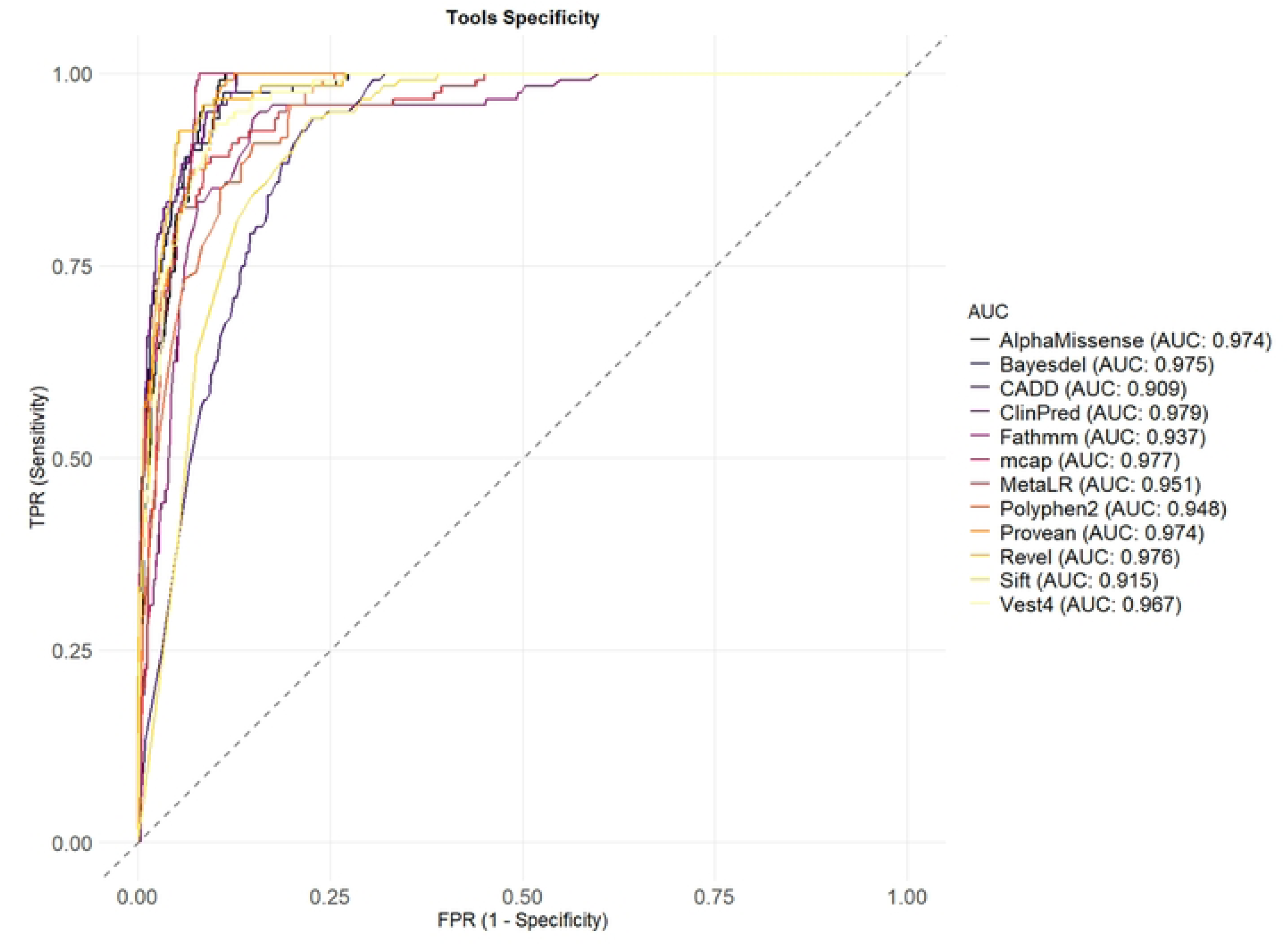
ROC curve displaying the accuracy of each nsSNP prediction model. TPR means True positive rate and FPR means False Positive Rate. FPR >0.5 for every tools.

### 3.3 Prediction and Analysis of Deleterious nsSNPs

A comprehensive computational approach employing twelve diverse in silico algorithms was utilized to accurately identify deleterious non-synonymous single-nucleotide polymorphisms (nsSNPs). These algorithms, including sequence homology-based tools like SIFT (score ≤ 0.05) and PROVEAN (score ≤-2.5), and those incorporating sequence and structural features such as PolyPhen-2 (score ≥ 0.85) and FATHMM (score < 0), were selected for their varied prediction mechanisms. Supervised machine learning models like AlphaMissense (score >0.5), VEST4 (score ≥ 0.5), M-CAP (score ≥ 0.025), and ClinPred (score ≥ 0.5) were also included, alongside ensemble meta predictors like MetaLR (score ≥ 0.5), BayesDel (score ≥ 0.2), REVEL (score ≥ 0.5), and CADD (PHRED score ≥ 20), which integrate multiple predictors for enhanced accuracy. Two datasets, comprising 2,583 nsSNPs in the TSC1 gene and 4,468 nsSNPs in the TSC2 gene, were analyzed. The integrated predictions from all twelve tools identified a total of 484 harmful nsSNPs in TSC1 and 1,268 harmful nsSNPs in TSC2. After a comprehensive in-silico analysis to get the most damaging nSNPs, a final set of twelve nsSNPs was identified as the most potentially damaging variants in TSC1 and sixteen in the TSC2 gene. These include rs1846545280, rs1846672141, rs2131961079, rs2132135077, rs2132135600, rs2132235475, rs2131703748, rs2132003126, rs2132003339, rs2132135678, rs2132135839, and rs2132154057 in TSC1 gene (Table 1).

**Table 1.**
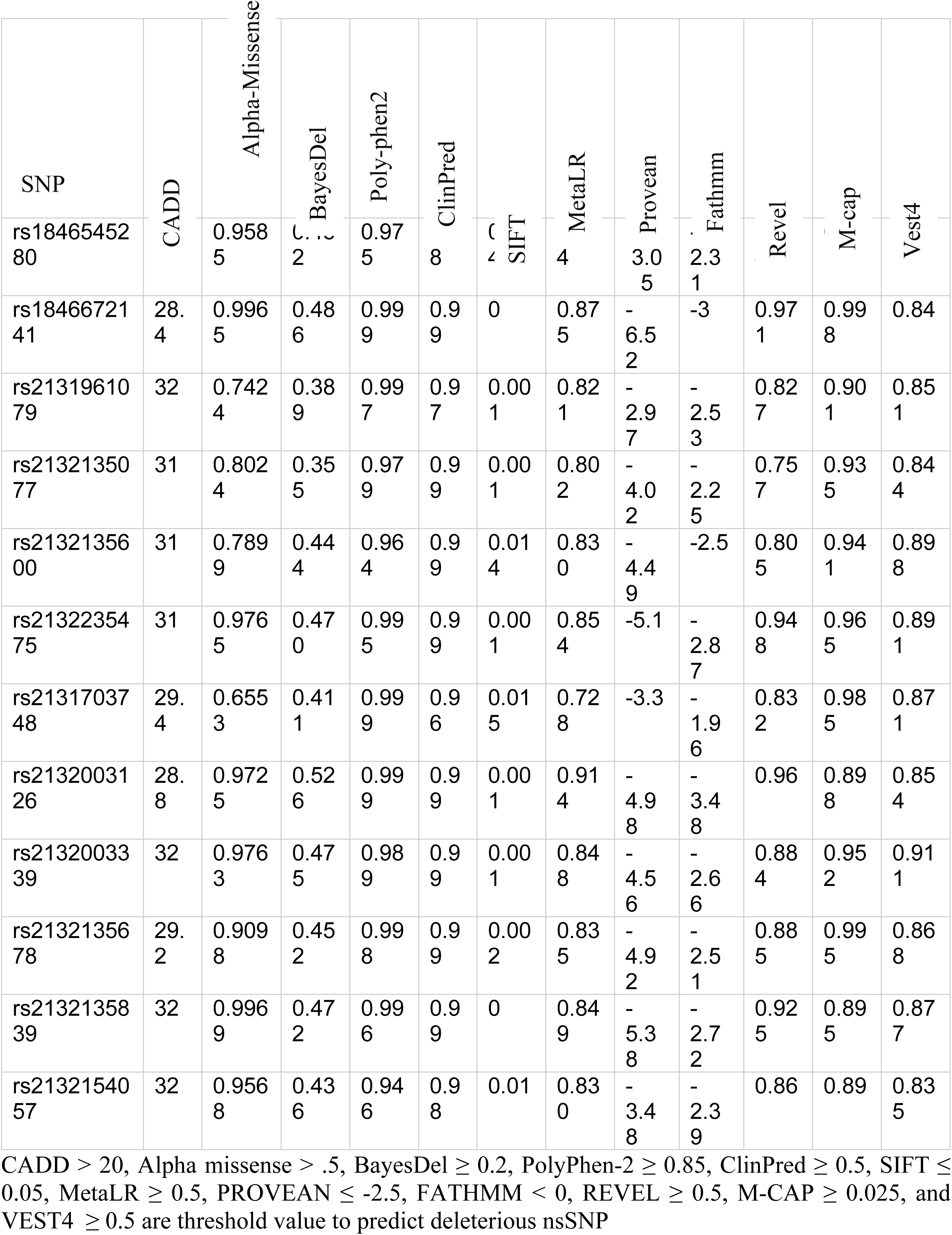
A compilation of 12 deleterious nsSNPs in the *TSC1* gene.

The nsSNPs in TSC2 are rs45517223, rs45517365, rs45517381, rs372918437, rs397514980, rs397515030, rs397515159, rs397515313, rs761588986, rs1163066622, rs1389060581, rs1555438434, rs1555438551, rs1555510327, rs2151354925, and rs2151550678 **(Table 2).** These variants consistently surpassed the deleterious prediction thresholds across all applied tools and remained as high confidence damaging nsSNPs after the final integrative analysis.

**Table 2.**
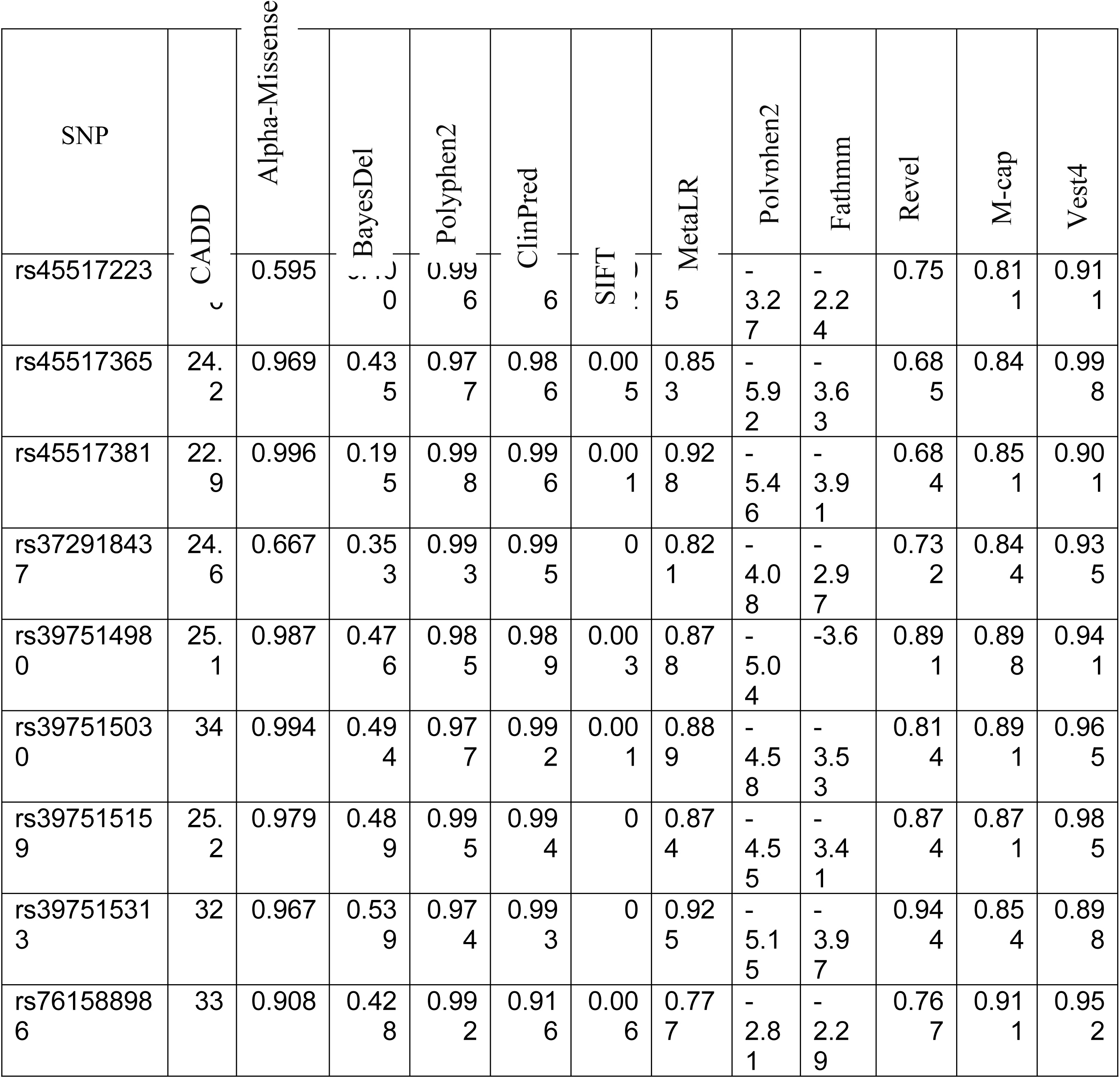
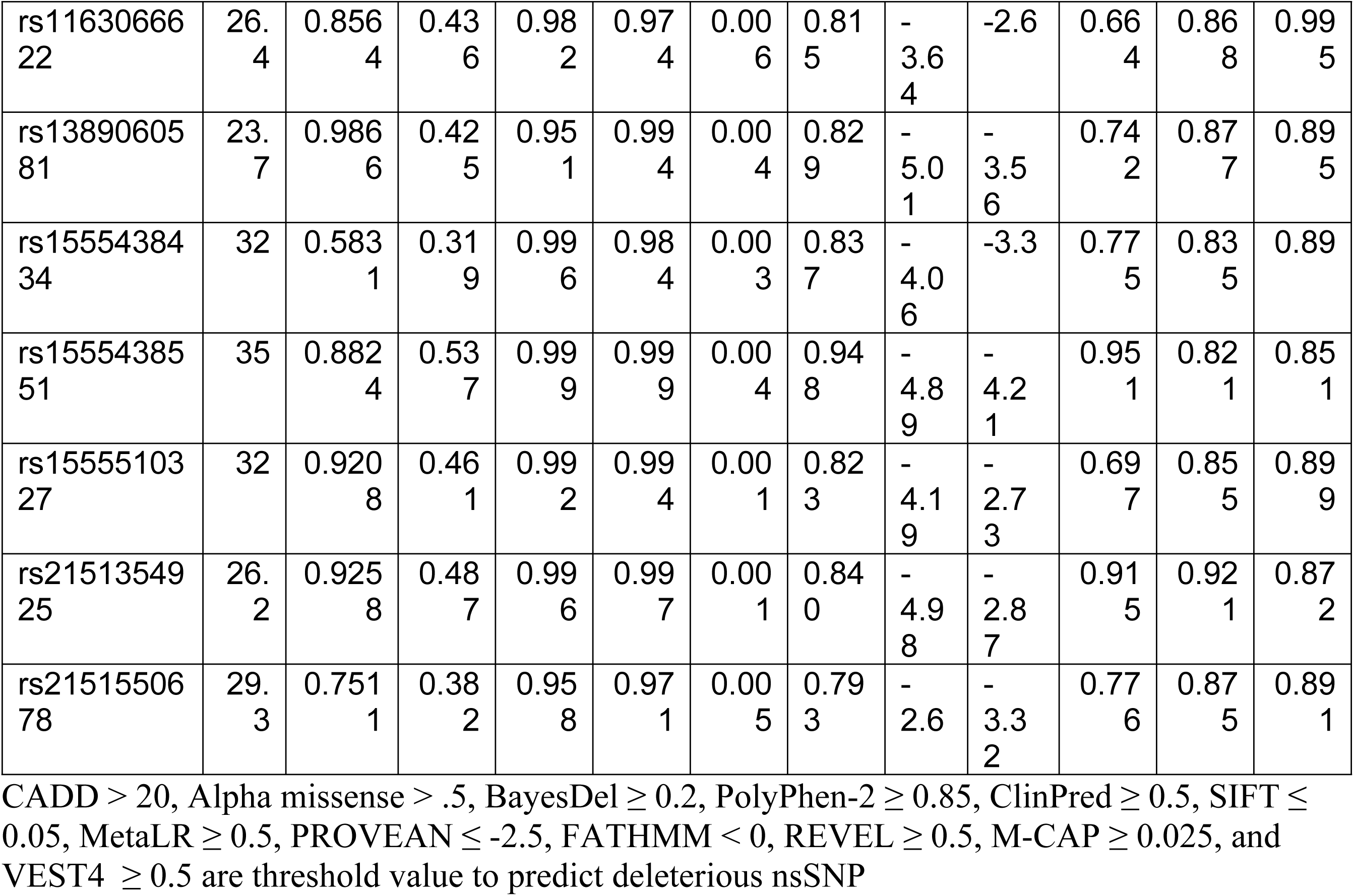
A compilation of sixteen deleterious nsSNPs in the *TSC2* gene.

### 3.4 Protein Stability Analysis

To evaluate protein stability, 484 and 1209 nsSNPs previously identified as harmful in TSC1 and TSC2, respectively, were further analyzed using two additional tools: I-Mutant 2.0 and MUpro. I-Mutant showed that 350 and 750 nsSNPs decreased protein stability in TSC1 and TSC2, respectively, using the reliability index (RI) value and changes in free energy (DDG).DDG values (kcal/mol) represent the change in Gibbs free energy upon mutation. Negative values (DDG < 0) suggest decreased protein stability, while positive values (DDG > 0) suggest increased stability. MUpro predicted that 300 nsSNPs destabilized the protein in TSC1, while 710 nsSNPs destabilized TSC2. For further analysis, missense variants of nsSNPs predicted to cause destabilization by both tools were selected. nsSNP selected by both tools are 250 in TSC1 and 650 in TSC2. Finally selected twelve nsSNPs of TSC1 and sixteen nsSNP of TSC2 are presented in Table 3 and Table 4.

**Table 3:** Impact of nsSNP on TSC1 protein stability.

| SNP | A.A.<br>position | I-mutant2.0 |  |  | MUpro |  |
| --- | --- | --- | --- | --- | --- | --- |
|  |  | Prediction | Reliability<br>index | DDG<br>Value(kcal/mol) | Prediction | DDG<br>Value(kcal/mol) |
| **rs1846545280** | G236E | Decrease | 7 | -1.19 | Decrease | -0.892 |
|  | G236A |  | 8 | -1.2 |  | -0.133 |
| **rs1846672141** | L161P | Decrease | 7 | -1.4 | Decrease | -1.51 |
| **rs2131850671** | I482K | Decrease | 7 | -0.7 | Decrease | -1.18 |
|  | I482R |  | 8 | -0.8 |  | -1.49 |
| **rs2131961079** | S361C | Decrease | 6 | -0.99 | Decrease | -1 |
|  | S361F |  | 7 | -0.91 |  | -1 |
| **rs2132135077** | S237Y | Decrease | 7 | -0.53 | Decrease | -1.32 |
|  | S237C |  | 6 | -0.91 |  | -1.27 |
|  | S237F |  | 6 | -0.67 |  | -1.51 |
| **rs2132135600** | T235S | Decrease | 8 | -2.31 | Decrease | -1.19 |
|  | T235I |  | 9 | -1.9 |  | -1.45 |
| **rs2132235475** | L90P | Decrease | 7 | -1.2 | Decrease | -1.21 |
| **rs2131703748** | L828H | Decrease | 8 | -1.15 | Decrease | -1.2 |
| **rs2132003126** | L264Q | Decrease | 8 | -1 | Decrease | -0.95 |
| **rs2132003339** | S263F | Increase | 3 | 1.1 | Decrease | -0.81 |
|  | S263C | Decrease | 8 | -1.5 |  | -0.62 |
| **rs2132135678** | V234E | Decrease | 7 | -0.98 | Decrease | -1.15 |
| **rs2132154057** | S201P | Increase | 4 | 1 | Increase | 0.7 |
|  | S201T | Decrease | 7 | -2.31 | Decrease | -1.5 |

DDG values (kcal/mol) represent the change in Gibbs free energy upon mutation. Negative values (DDG < 0) suggest decreased protein stability, while positive values (DDG > 0) suggest increased stability

**Table 4:**
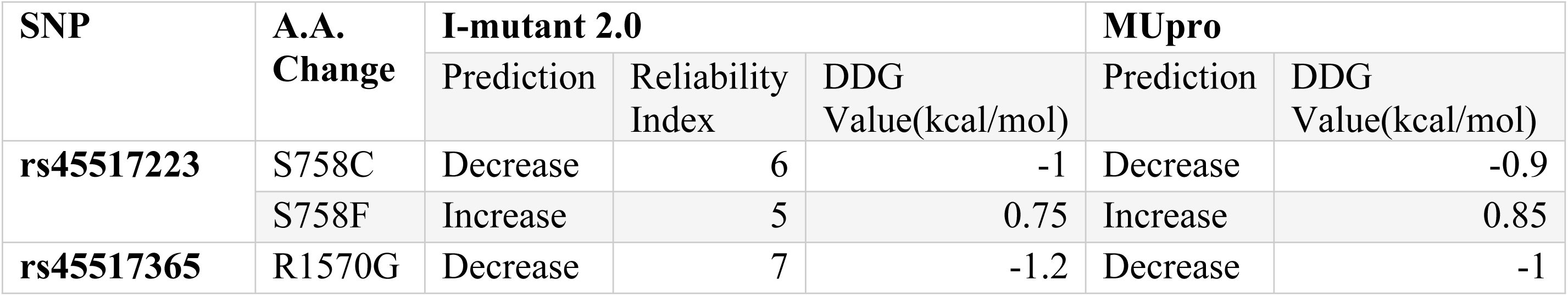
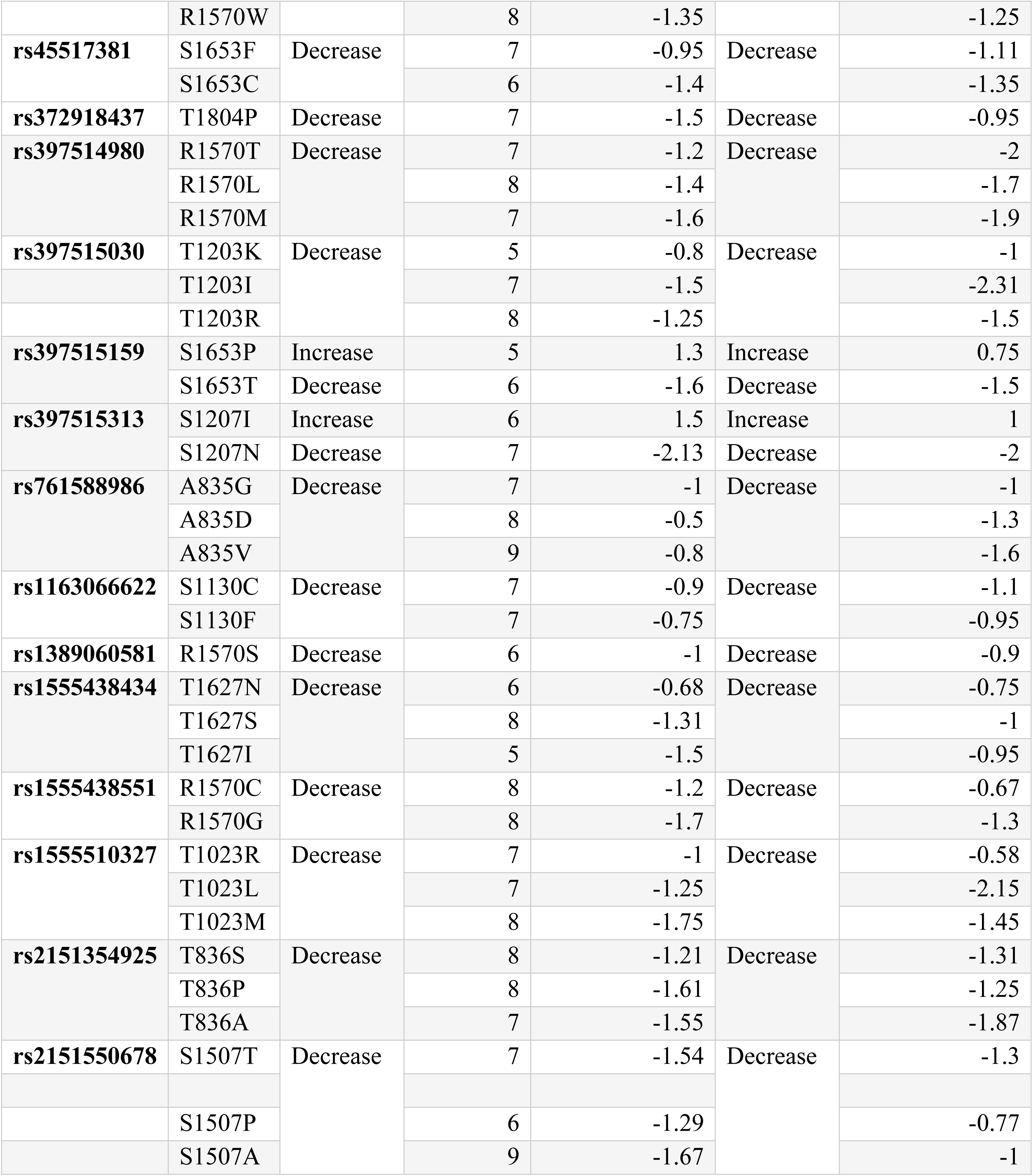
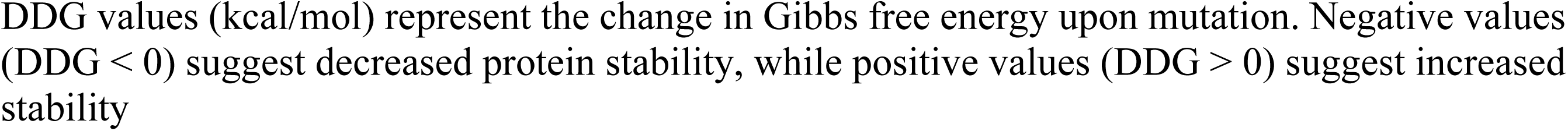
Impact of nsSNP on TSC2 protein stability.

| SNP | A.A.<br>Change | I-mutant 2.0 |  |  | MUpro |  |
| --- | --- | --- | --- | --- | --- | --- |
|  |  | Prediction | Reliability<br>Index | DDG<br>Value(kcal/mol) | Prediction | DDG<br>Value(kcal/mol) |
| **rs45517223** | S758C | Decrease | 6 | -1 | Decrease | -0.9 |
|  | S758F | Increase | 5 | 0.75 | Increase | 0.85 |
| **rs45517365** | R1570G | Decrease | 7 | -1.2 | Decrease | -1 |
|  | R1570W |  | 8 | -1.35 |  | -1.25 |
| **rs45517381** | S1653F | Decrease | 7 | -0.95 | Decrease | -1.11 |
|  | S1653C |  | 6 | -1.4 |  | -1.35 |
| **rs372918437** | T1804P | Decrease | 7 | -1.5 | Decrease | -0.95 |
| **rs397514980** | R1570T | Decrease | 7 | -1.2 | Decrease | -2 |
|  | R1570L |  | 8 | -1.4 |  | -1.7 |
|  | R1570M |  | 7 | -1.6 |  | -1.9 |
| **rs397515030** | T1203K | Decrease | 5 | -0.8 | Decrease | -1 |
|  | T1203I |  | 7 | -1.5 |  | -2.31 |
|  | T1203R |  | 8 | -1.25 |  | -1.5 |
| **rs397515159** | S1653P | Increase | 5 | 1.3 | Increase | 0.75 |
|  | S1653T | Decrease | 6 | -1.6 | Decrease | -1.5 |
| **rs397515313** | S1207I | Increase | 6 | 1.5 | Increase | 1 |
|  | S1207N | Decrease | 7 | -2.13 | Decrease | -2 |
| **rs761588986** | A835G | Decrease | 7 | -1 | Decrease | -1 |
|  | A835D |  | 8 | -0.5 |  | -1.3 |
|  | A835V |  | 9 | -0.8 |  | -1.6 |
| **rs1163066622** | S1130C | Decrease | 7 | -0.9 | Decrease | -1.1 |
|  | S1130F |  | 7 | -0.75 |  | -0.95 |
| **rs1389060581** | R1570S | Decrease | 6 | -1 | Decrease | -0.9 |
| **rs1555438434** | T1627N | Decrease | 6 | -0.68 | Decrease | -0.75 |
|  | T1627S |  | 8 | -1.31 |  | -1 |
|  | T1627I |  | 5 | -1.5 |  | -0.95 |
| **rs1555438551** | R1570C | Decrease | 8 | -1.2 | Decrease | -0.67 |
|  | R1570G |  | 8 | -1.7 |  | -1.3 |
| **rs1555510327** | T1023R | Decrease | 7 | -1 | Decrease | -0.58 |
|  | T1023L |  | 7 | -1.25 |  | -2.15 |
|  | T1023M |  | 8 | -1.75 |  | -1.45 |
| **rs2151354925** | T836S | Decrease | 8 | -1.21 | Decrease | -1.31 |
|  | T836P |  | 8 | -1.61 |  | -1.25 |
|  | T836A |  | 7 | -1.55 |  | -1.87 |
| **rs2151550678** | S1507T | Decrease | 7 | -1.54 | Decrease | -1.3 |
|  | S1507P |  | 6 | -1.29 |  | -0.77 |
|  | S1507A |  | 9 | -1.67 |  | -1 |

DDG values (kcal/mol) represent the change in Gibbs free energy upon mutation. Negative values (DDG < 0) suggest decreased protein stability, while positive values (DDG > 0) suggest increased stability

### 3.5 Evolutionary Conservation Analysis of High-Risk nsSNPs

Highly conserved regions hold critical roles in maintaining the molecule’s function and structure, where changes are often deleterious, while more variable regions tolerate mutations without significantly compromising the macromolecule’s performance (52).ConSurf analysis revealed that 20 nsSNPs are located within highly conserved region and the rest in variable regions. Only SNPs in highly conserved regions were selected for further analysis. Among the final twelve nsSNPs of TSC1, G236, S361, T235, and S263 are considered functional, highly conserved and exposed residues. L161, I482, S237, L90, L828, L264, V234, and S201 are considered highly conserved, structural, and buried residues **(Table 5)**.

**Table 5:**
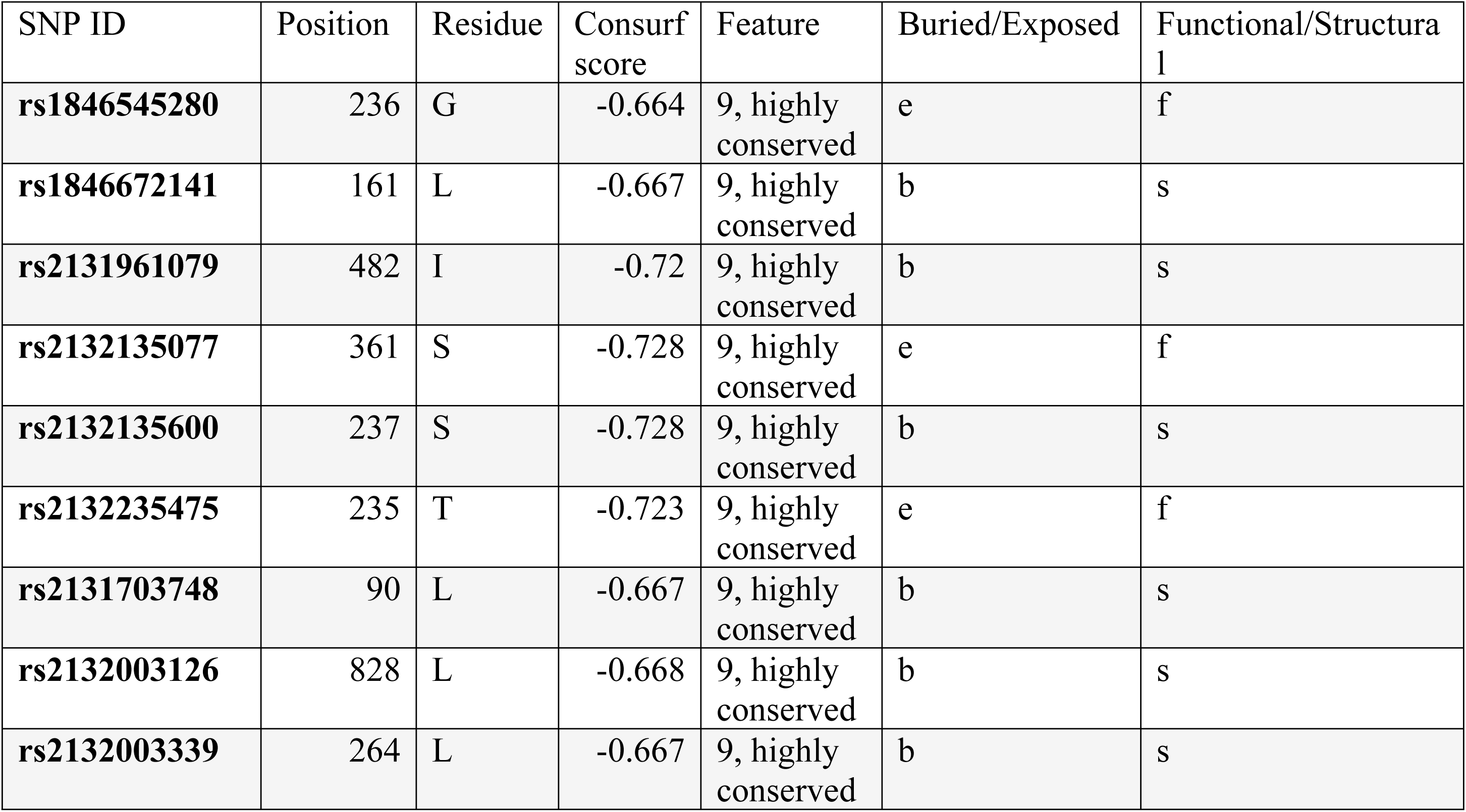
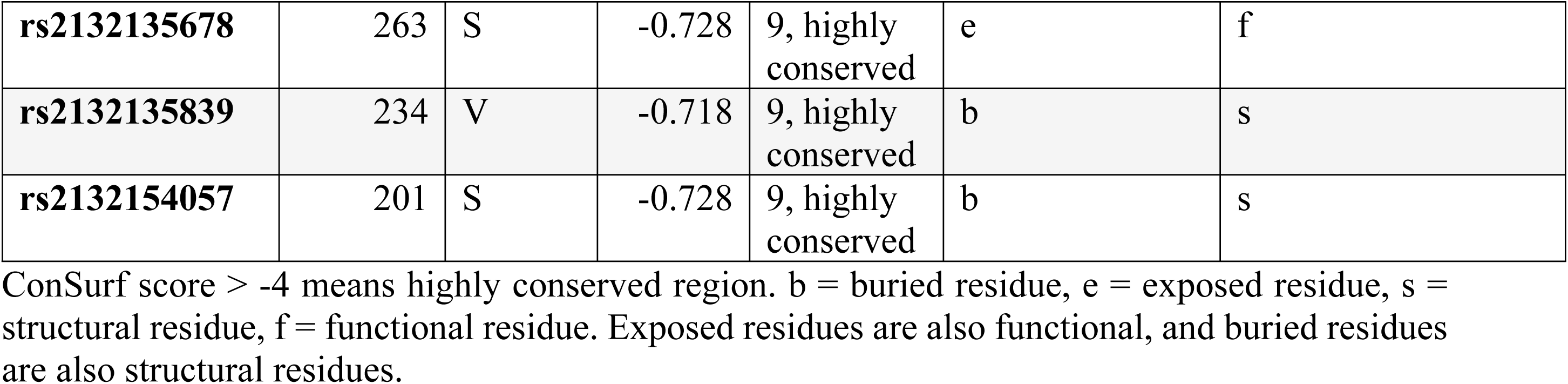
Conservation score of nsSNPs in *TSC1*.

| SNP ID | Position | Residue | Consurf score | Feature | Buried/Exposed | Functional/Structural |
| --- | --- | --- | --- | --- | --- | --- |
| **rs1846545280** | 236 | G | -0.664 | 9, highly conserved | e | f |
| **rs1846672141** | 161 | L | -0.667 | 9, highly conserved | b | s |
| **rs2131961079** | 482 | I | -0.72 | 9, highly conserved | b | s |
| **rs2132135077** | 361 | S | -0.728 | 9, highly conserved | e | f |
| **rs2132135600** | 237 | S | -0.728 | 9, highly conserved | b | s |
| **rs2132235475** | 235 | T | -0.723 | 9, highly conserved | e | f |
| **rs2131703748** | 90 | L | -0.667 | 9, highly conserved | b | s |
| **rs2132003126** | 828 | L | -0.668 | 9, highly conserved | b | s |
| **rs2132003339** | 264 | L | -0.667 | 9, highly conserved | b | s |
| **rs2132135678** | 263 | S | -0.728 | 9, highly conserved | e | f |
| **rs2132135839** | 234 | V | -0.718 | 9, highly conserved | b | s |
| **rs2132154057** | 201 | S | -0.728 | 9, highly conserved | b | s |

ConSurf score > -4 means highly conserved region. b = buried residue, e = exposed residue, s = structural residue, f = functional residue. Exposed residues are also functional, and buried residues are also structural residues.

In case of TSC2, ConSurf shows that 27 nsSNPs are in highly conserved regions. Among the final sixteen nsSNPs of *TSC2*, S758, R1570, S1653, T1804, S1203, T1627, T1023) are revealed as exposed, highly conserved and functional residues. S1207, A835, S1130, T836, and S1507 are buried, highly conserved, and structural (Table 6).

**Table 6:** Conservation score output of nsSNPs in *TSC2*.

| SNP ID | Position | Residue | ConSurf score | Feature | Exposed/Buried | Structural/Functional |
| --- | --- | --- | --- | --- | --- | --- |
| **rs45517223** | 758 | S | -0.512 | 9, highly conserved | e | f |
| **rs45517365** | 1570 | R | -0.492 | 9, highly conserved | e | f |
| **rs45517381** | 1653 | S | -0.512 | 9, highly conserved | e | f |
| **rs372918437** | 1804 | T | -0.508 | 9, highly conserved | e | f |
| **rs397514980** | 1570 | R | -0.492 | 9, highly conserved | e | f |
| **rs397515030** | 1203 | T | -0.508 | 9, highly conserved | e | f |
| **rs397515159** | 1653 | S | -0.512 | 9, highly conserved | e | f |
| **rs397515313** | 1207 | S | -0.512 | 9, highly conserved | b | s |
| **rs761588986** | 835 | A | -0.504 | 9, highly conserved | b | s |
| **rs1163066622** | 1130 | S | -0.512 | 9, highly conserved | b | s |
| **rs1389060581** | 1570 | R | -0.492 | 9, highly conserved | e | f |
| **rs1555438434** | 1627 | T | -0.508 | 9, highly conserved | e | f |
| **rs1555438551** | 1570 | R | -0.492 | 9, highly conserved | e | f |
| **rs1555510327** | 1023 | T | -0.508 | 9, highly conserved | e | f |
| **rs2151354925** | 836 | T | -0.508 | 9, highly conserved | b | s |
| **rs2151550678** | 1507 | S | -0.512 | 9, highly conserved | b | s |

ConSurf score > -4 means highly conserved region. b = buried residue, e = exposed residue, s = structural residue, f = functional residue. Exposed residues are also functional, and buried residues are also structural residues.

### 3.6 Prediction of Post-Translation Modification (PTM) Site

NetPhos 3.1 was employed to get phosphorylation sites in both *TSC1* and *TSC2*. The threshold value for any amino acid position to be predicted as a phosphorylation site is >0.5. Significant Phosphorylation sites of TSC1 and TSC2 are shown in **Fig 3 and Fig 4** respectively. Result table is given in supporting table **S1**.

**Fig 3:**
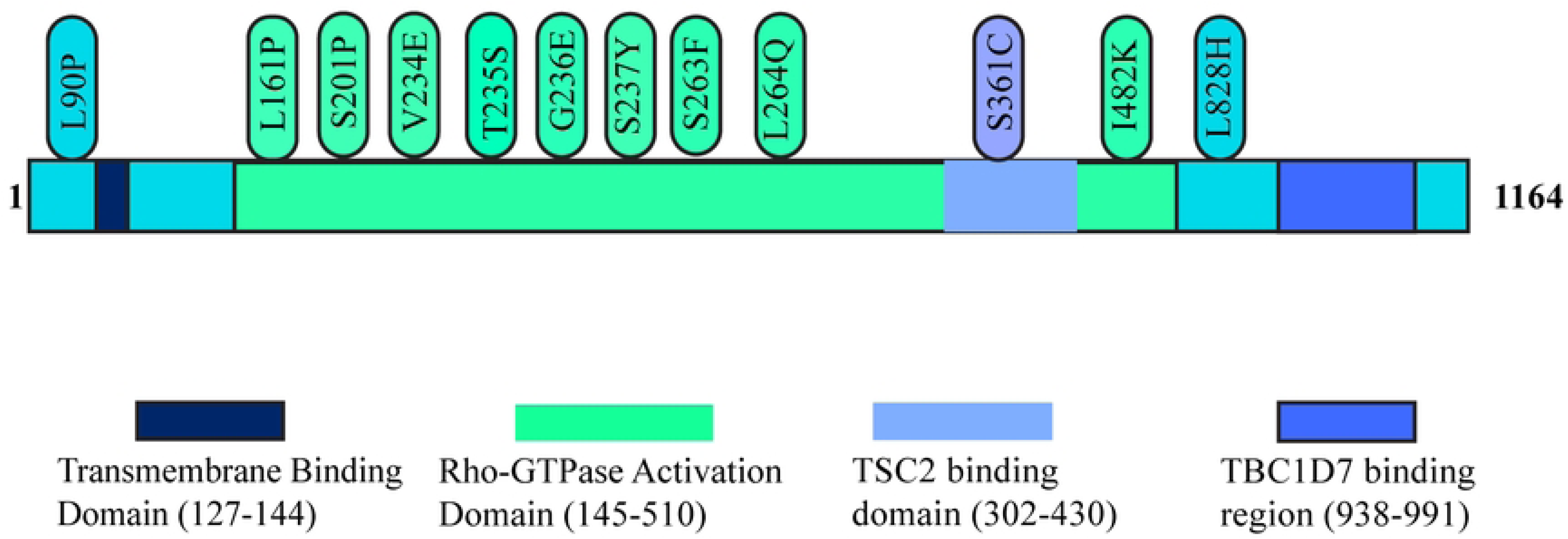
Predicted Phosphorylation site in *TSC1*. S201P, T235S, S237Y, S263F, G236E, L161P, S237Y, L264Q, V234E, I482K are in Rho-GTPase activating domain. S361C is in the TSC2 binding domain. L828H is in the NFL binding domain.

**Fig 4:**
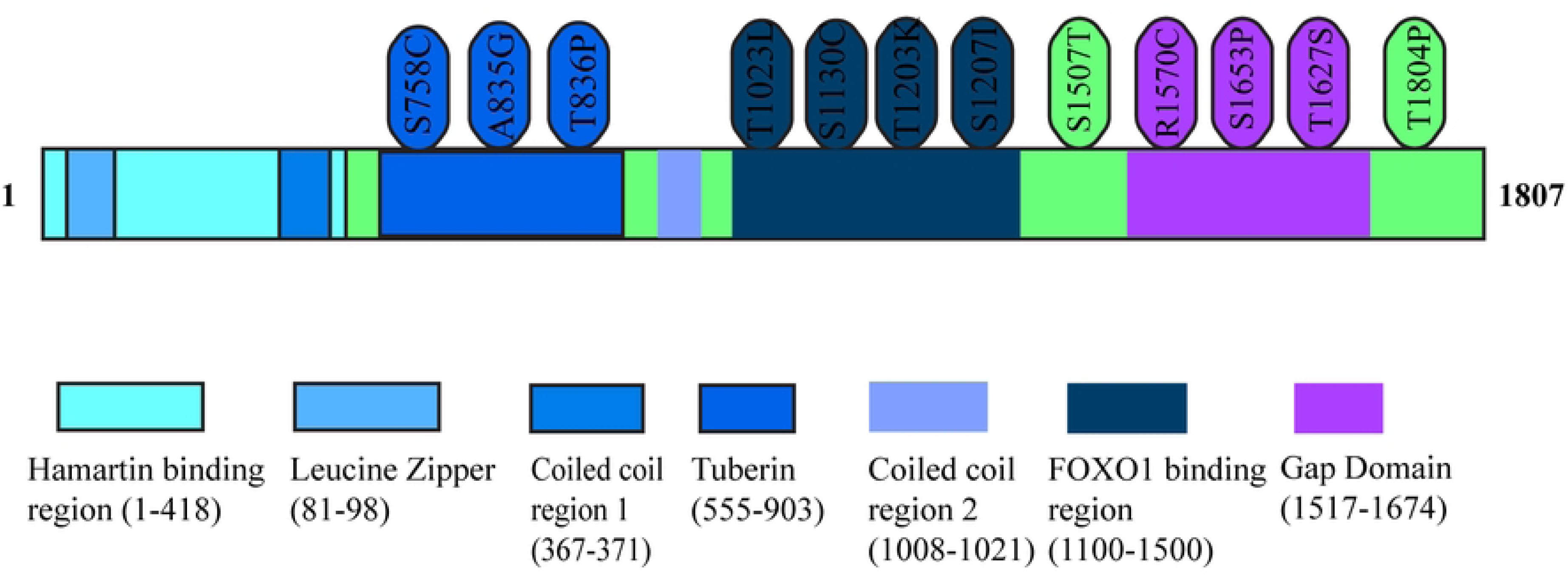
Predicted Phosphorylation site in TSC2. S758C, A835G, and T836P are in the tuberin region. T1023L, S1130C, T1203K, S1207I, and S1507T are present on FOXO1 binding region. R1570C, T1627S, S1653P, and T1804P are in the GAP domain

### 3.7 Identification of Functional and Structural Modifications

The 12 most damaging SNPs of TSC1 and 16 of TSC2 were submitted to the MutPred web server and the findings with probability scores are given in **Table 7** and **Table 8**, respectively. All nsSNPs show strong pathogenic potential with g value above 0.6 and p-value less than 0.05.

**Table 7:** MutPred2 prediction of the TSC1 nsSNPs.

| Mutation | Actionable/confident hypothesis | Mutpred Score | Probability | P value |
| --- | --- | --- | --- | --- |
| **G236E** | Gain of intrinsic disorder | 0.824 | 0.38 | 0.01 |
|  | Loss of strand |  | 0.27 | 0.02 |
|  | Altered Metal binding |  | 0.15 | 0.04 |
| **G236A** | Gain of Helix | 0.662 | 0.27 | 0.04 |
|  | Loss of strand |  | 0.27 | 0.03 |
|  | Altered Metal binding |  | 0.14 | 0.05 |
|  | Altered Stability |  | 0.11 | 0.04 |
| **L161P** | Gain of Allosteric site at R160 | 0.936 | 0.19 | 0.05 |
| **I482K** | Gain of B-factor | 0.782 | 0.27 | 0.01 |
| **I482R** | Gain of SUMOylation at K477 |  | 0.26 | 0.00072 |
|  | Altered Coiled coil |  | 0.21 | 0.02 |
|  | Gain of Ubiquitylation at I482 |  | 0.21 | 0.00053 |
|  | Gain of Methylation at I482 |  | 0.1 | 0.04 |
| **S237Y** | Loss of Intrinsic order | 0.78 | 0.35 | 0.04 |
|  | Gain of Allosteric site at E421 |  | 0.19 | 0.04 |
|  | Altered Metal binding |  | 0.14 | 0.04 |
|  | Gain of Sulfation at S237 |  | 0.02 | 0.02 |
| **S237C** | Loss of Intrinsic order | 0.688 | 0.36 | 0.04 |
|  | Altered Metal binding |  | 0.32 | 0.01 |
|  | Loss of Strand |  | 0.28 | 0.00078 |
| **S237F** | Loss of Intrinsic order | 0.754 | 0.36 | 0.04 |
|  | Gain of Acetylation at K238 |  | 0.2 | 0.04 |
|  | Altered Metal binding |  | 0.16 | 0.03 |
| **T235I** | Loss of Strand | 0.705 | 0.26 | 0.04 |
|  | Altered Metal binding |  | 0.25 | 0.04 |
| **L90P** | Altered Stability | 0.906 | 0.09 | 0.05 |
| **L828H** | Gain of Intrinsic disorder | 0.727 | 0.33 | 0.03 |
|  | Altered Coiled coil |  | 0.14 | 0.03 |
|  | Altered Stability |  | 0.13 | 0.03 |
| **L264Q** | Altered Metal binding | 0.883 | 0.27 | 0.02 |
|  | Altered Stability |  | 0.18 | 0.01 |
|  | Altered Transmembrane protein |  | 0.17 | 0.01 |
| **S263F** | Altered Metal binding | 0.773 | 0.36 | 0.00084 |
|  | Altered Transmembrane protein |  | 0.14 | 0.02 |
| **S263C** | Altered Metal binding | 0.757 | 0.25 | 0.03 |
|  | Altered Transmembrane protein |  | 0.14 | 0.02 |
| **V234E** | Gain of Intrinsic disorder | 0.918 | 0.47 | 0.00023 |
|  | Loss of Strand |  | 0.27 | 0.02 |
|  | Altered Metal binding |  | 0.25 | 0.03 |
|  | Altered Stability |  | 0.14 | 0.02 |

MutPred score >6 indicates potential pathogenicity. Probability means confidence in prediction. P-value <0.05 means significance.

**Table 8:** MutPred2 prediction of TSC2 nsSNPs.

| **Mutation** | **Actionable/confident hypothesis** | **Mutpred score** | **Probability** | **P value** |
| --- | --- | --- | --- | --- |
| **S1653F** | Altered Ordered interface, metal binding, Transmembrane protein | 0.885 | 0.35,0.26,0.21 | 0.00045, 0.03, 0.00039 |
|  | Loss of Relative solvent accessibility |  | 0.3 | 0.00094 |
|  | Loss of Allosteric site at Y1650, loss of sulfation |  | 0.23, 0.02 | 0.02, 0.04 |
|  | Loss of N-linked glycosylation at N1651 |  | 0.12 | 0.00098 |
| **S1653C** | Gain of Allosteric site at Y1650 | 0.842 | 0.25 | 0.01 |
| **T1804P** | Gain of Intrinsic disorder | 0.875 | 0.33, 0.25 | 0.03, 0.03 |
|  | Altered Disordered interface, Altered metal binding |  | 0.28, 0.25 | 0.04, 0.03 |
|  | Loss of Strand, Loss of Relative solvent accessibility |  | 0.26, 0.25 | 0.04 |
| **T1203K** | Gain of Allosteric site at W1208, Gain of N-linked glycosylation at N1205 | 0.797 | 0.29, 0.04 | 0.0027, 0.2 |
|  | Altered Transmembrane protein, Altered DNA binding |  | 0.19, 0.14 | 0.0063, 0.05 |
| **T1205I** | Altered Ordered interface, Transmembrane protein | 0.752 | 0.24, 0.18 | 0.04, 0.0087 |
| **T1203R** | Loss of N-linked glycosylation at N1205 | 0.854 | 0.04 | 0.03 |
| **S1653P** | Altered Metal binding, ordered interface, Transmembrane protein | 0.93 | 0.32, 0.03, 0.21 | 0.01, 0.02, 0.0036 |
|  | Loss of Relative solvent accessibility, N-linked glycosylation at N1651, Sulfation at Y1650 |  | 0.31, 0.12, 0.02 | 0.0074, 0.0098, 0.04 |
|  | Gain of Allosteric site at Y1650 |  | 0.25 | 0.01 |
| **S1653T** | Gain of N-linked glycosylation at N1651 |  | 0.14 | 0.0064 |
| **S1207I** | Altered Ordered interface, Transmembrane protein | 0.914 | 0.27, 0.17 | 0.0093, 0.0094 |
|  | Loss of Allosteric site at W1208, N-linked glycosylation at N1205 |  | 0.23, 0.04 | 0.02, 0.01 |
| **S1207N** | Gain of Allosteric site at W1208 | 0.877 | 0.25 | 0.01 |
| **S1130F** | Loss of Phosphorylation at S1130 | 0.634 | 0.57 | 0.0026 |
|  | Altered Disordered interface |  | 0.27 | 0.01 |
| **T1023L** | Altered Transmembrane protein | 0.872 | 0.16 | 0.01 |
| **T836P** | Altered Transmembrane protein | 0.806 | 0.12 | 0.03 |
| **S1507P** | Loss of Loop | 0.829 | 0.28 | 0.02 |
| **R1570G/W** | Altered Ordered interface, Transmembrane protein | 0.799,0.777 | 0.25, 0.29 | 0.03, 0.05 |
| **R1570T/L/M** | Altered Ordered interface, Transmembrane protein | 0.8,0.799, 0.734 | 0.25, 0.11 | 0.03, 0.04 |
| **A835D** | Altered Transmembrane protein | 0.81 | 0.16 | 0.04 |
| **R1570S** | Gain of Relative solvent accessibility | 0.767 | 0.25 | 0.03 |
|  | Altered Ordered interface, Transmembrane protein |  | 0.24, 0.1 | 0.05, 0.03 |
| **R1570C** | Altered Ordered interface, Transmembrane protein | 0.764 | 0.24, 0.9 | 0.04,0.05 |
|  | Loss of Relative solvent accessibility |  | 0.24 | 0.05 |

MutPred score >6 indicates potential pathogenicity. Probability means confidence in prediction. P-value <0.05 means significance.

### 3.8 Estimating the Effects of High Risk nsSNPs on the Protein Structure

All twelve nsSNPs resulted in changes in amino acid size. Three mutant amino acids (G236E/A, I482K/R, V234E) exhibited both increased size and less hydrophobicity compared to their wild-type counterparts in *TSC1*. Five nsSNPs (S237C/Y/F, T235I/S, L828H, L264Q, S263C/F) were found to be both of increased size and more hydrophobic. Two (L161P, L90P) nsSNPs are smaller in size and neutral in charge **(Table 9)**.

**Table 9:** Project HOPE analysis of TSC1.

| SNP ID | A.A | Aminoacid properties | Impact on *TSC1* |
| --- | --- | --- | --- |
| rs1846672141 | L161P | Mutant residue is smaller in size. | Might result in loss of interaction. |
| rs2132235475 | L90P | The mutant residue is smaller and both are neutral. | Proline disrupts an α-helix when not located at one of the first 3 positions of that helix; thus, the helix will be disturbed. |
| rs1846545280 | G236E/A | The mutant residue is negative, bigger and less hydrophobic. Only glycine is flexible enough to make the necessary torsion angles, mutation may disrupt local backbone into an incorrect conformation. | The wild-type residue is a glycine, the most flexible of all residues. This flexibility might be necessary for the protein's function. Mutation of this glycine can abolish this function. |
| rs2131850671 | I482K/R | A mutant residue is larger and less hydrophobic, which may lead to the loss of hydrophobic interactions at the surface or core. | The mutation is within a special region that mediates interaction with WDR45B, thus affecting the interaction. |
| rs2132135077 | S237C/Y/F | Mutated residues are bigger and more hydrophobic, which may result in loss of a hydrogen bond. | The wild-type residue is predicted to be in a turn, while the mutant residue is in another secondary structure; therefore, the local conformation will be slightly destabilized. |
| rs2132135600 | T235I/S | Mutated residue is bigger and more hydrophobic. The mutant residue is smaller, this might lead to loss of interactions. | Isoleucine causes a bump and can lead to loss of hydrogen bonds. The wild-type residue is in β-strand. Serine prefers to be in another secondary structure; thus, the local conformation will be slightly destabilized. |
| rs2131703748 | L828H | The mutated residue is bigger and more hydrophobic so it may lead to a bump. | The wild-type residue is in an α-helix, while the mutated residue does not prefer α-helices. |
| rs2132003126 | L264Q | The mutated residue is bigger and more hydrophobic so it may lead to a bump. | Hydrophobic interactions, either in the core of the protein or on the surface, will be lost due to mutation. |
| rs2132003339 | S263C/F | Alanine is a more hydrophobic residue, whereas phenylalanine is both bigger and more hydrophobic, leading to a bump. | A mutation may cause loss of hydrogen bonds. |
| rs2132135678 | V234E | Mutant residue is bigger, less hydrophobic, leading to loss of hydrophobic interaction, and negatively charged. | The wild-type residue is predicted to be located in a β-strand. The mutant residue prefers to be in another secondary structure; thus the local conformation will be slightly destabilized. |
| rs2132154057 | S201P/T | Proline is bigger and more hydrophobic. Threonine is larger. | This mutation can lead to loss of a hydrogen bond. Threonine can lead to bump. |

The difference between properties of wild-type and mutant-type amino acids regarding size, charge, and hydrophobicity is displayed.

Nine nsSNPs (S758C/F, S1653C/F, S1653T/P, T1207N/I, S1130C/F, T1627N/S/I, T1023L/M, A835D/G/V, S1507T/P/A) were bigger and more hydrophobic than the wild type. T1203R/K/I has positively charged mutated residues. Both T836P/A and T1804P got small and more hydrophobic mutated residues. In R1570G/W, R1570T/L/M and R1570S, arginine is replaced by small, neutral, and more hydrophobic residues. A835D/G/V has small but more hydrophobic mutated residues (Table 10).

**Table 10:** Project HOPE analysis of *TSC2*.

| SNP ID | A. A | Amino acid properties | Impact on *TSC2* |
| --- | --- | --- | --- |
| rs45517223 | S758C/F | The mutation introduces a more hydrophobic residue, resulting in loss of hydrogen bonds. Wild-type residue is 100% conserved. | The wild-type residue is in its preferred secondary structure, a turn. The mutant residue prefers to be in another secondary structure; so, the local conformation will be slightly destabilized. |
| rs45517381 | S1653C | Mutation introduces a more hydrophobic residue in place of a 100% conserved wild-type residue. | The wild-type residue forms a hydrogen bond with 1651 and 1655. The mutation will cause loss of hydrogen bonds in the core of the protein and, as a result, disturb correct folding. |
|  | S1653F | The wild-type residue was buried in the core of the protein. The mutant residue is bigger and more hydrophobic. | The mutated residue is located in a domain that is important for the activity of the protein and in contact with residues in another domain. Mutation can cause an altered reaction. |
| rs372918437 | T1804P | The mutant residue is more hydrophobic than the wild-type residue. | The mutation occurs in place of a 100% conserved residue, damaging the protein. Also, it disrupts the hydrogen bond. |
| rs397515030 | T1203R/K/I | Mutant residue is positively charged, bigger, and more hydrophobic. | Hydrophobic reaction at the core or surface in tuberin domain may be lost and it also may lead to bump. |
| rs397515159 | S1653T/P | Threonine is bigger while proline is both bigger and hydrophobic. | A large and hydrophobic mutated residue disrupts interactions within the Rap-Gap domain. Proline, being a hydrophobic residue, will cause loss of hydrogen bond. |
| rs397515313 | S1207N/I | Mutant residue is bigger and more hydrophobic. | Asparagine being the more hydrophobic one, hydrophobic interactions, either in the core of the protein or on the surface, will be lost. A bigger size of the mutant may lead to a bump and hydrophobicity will cause loss of hydrogen bond. |
| rs1163066622 | S1130C/F | Mutated residue is big, hydrophobic and in place of a 100% conserved residue, | Mutation is in disordered region, and due to increased hydrophobicity, it will cause loss of hydrogen bonds. |
| rs1555438434 | T1627N/S/I | Both Asparagine and Isoleucine are more bigger than serine and hydrophobic residue. | It may lead to loss of interaction with other molecules on surface and also affect hydrophobic interaction. Serine is smaller residue which may cause loss of external interaction. |
| rs1555510327 | T1023L/M | Mutant residue is bigger and more hydrophobic. | Bigger residue can cause bump. Also, loss of hydrogen bond is possible. |
| rs2151354925 | T836P/A/S | Alanine and serine are both small and hydrophobic. Proline is hydrophobic. | Introduction of a more hydrophobic residue in place of a conserved residue in the tuberin domain may lead to the loss of a hydrogen bond. |
| rs2151550678 | S1507P/T/A | Proline is bigger and more hydrophobic. Threonine is also bigger which can cause bump. Alanine is small and more hydrophobic residue. | Serine is in a turn whereas mutated residues are preferably in another secondary structure. Presence of bigger and more hydrophobic residue in Gap domain can cause loss of hydrogen bond thus affecting protein folding. |
| rs45517365 | R1570G/W | Mutant residue is neutral. Arginine form salt bridge among 1652, 1677 and 1705 | Glycine can disturb the required rigidity of the protein at this position. Bigger residue leads to loss of interaction with other residues in the Gap domain. |
| rs397514980 | R1570T/L/M | The mutant residue is smaller and more hydrophobic. | A mutation may lead to loss of external interaction and also loss of interaction with other molecules. |
| rs761588986 | A835D/G/V | Aspartic acid is bigger, more hydrophobic, and negatively charged. Glycine is smaller and more hydrophobic. Valine is bigger. This position is 100% conserved. | Mutation to aspartic acid may lead to loss of reaction at the core or on the surface and repulse the same charged residues. Glycine can disturb the required rigidity of the protein at this position and also cause loss of interaction. Valine may lead to a bump due to its size. |
| rs1389060581 | R1570S | The mutant residue is smaller than the wild-type residue. | This will cause a possible loss of external interactions. |

The difference between properties of wild-type and mutant-type amino acids regarding size, charge, and hydrophobicity is showed.

### 3.9 Functional Enrichment and PPI Network Analysis

Functional enrichment analysis was conducted with KEGG and DisGenet. KEGG pathway shows significantly enriched pathway due to TSC1 and TSC2 **(Fig 5 A)**. Concurrently, DisGeNET reveals significantly enriched diseases with both genes **(Fig 5 B)**. An analysis of the TSC1 and TSC2 proteins’ interaction network using the STRING server revealed interactions with 11 genes **(Fig 6)**.

**Fig 5:**
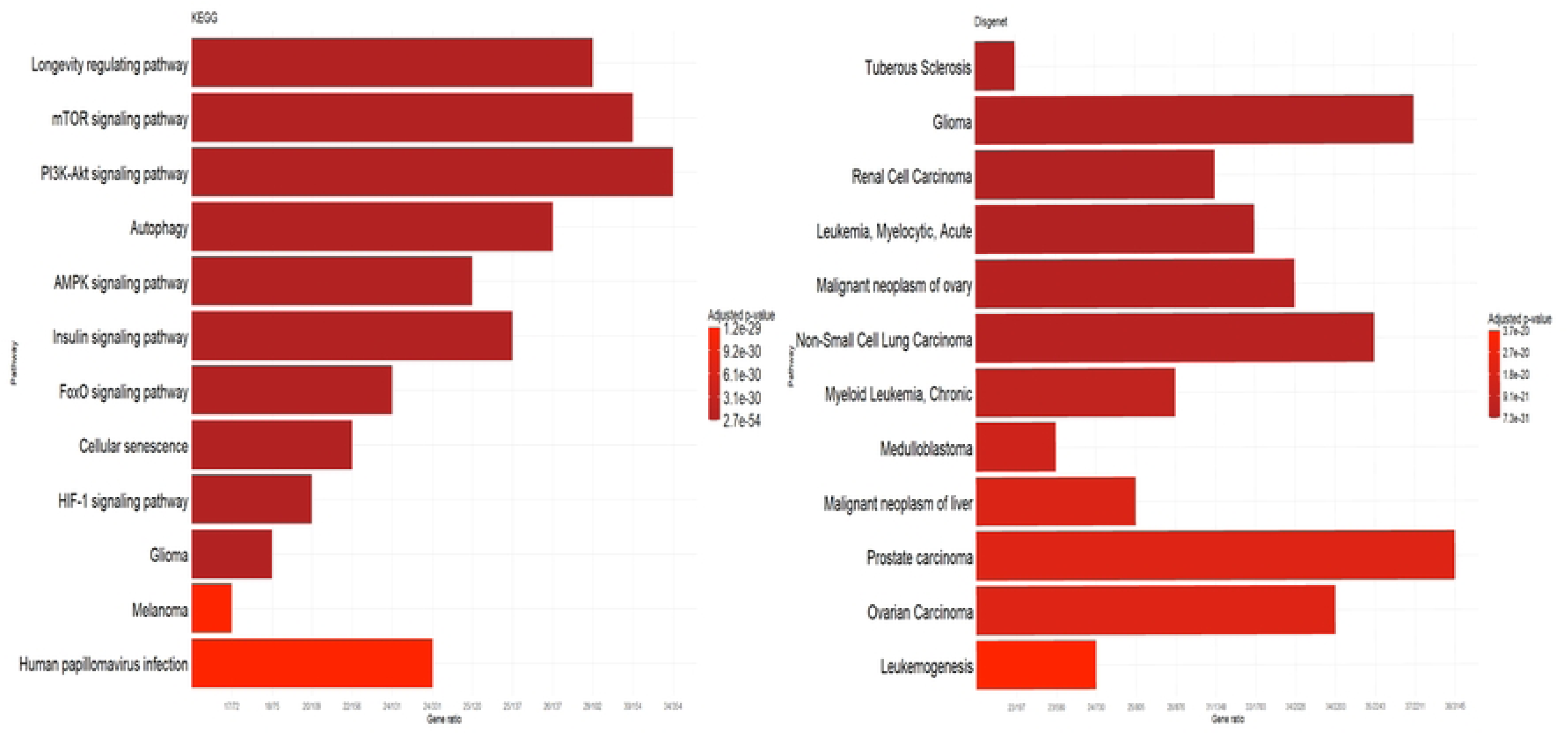
Functional enrichment analysis in biological pathways. (A) KEGG pathways display associations of both TSC1 and TSC2 with different pathways. (B) DisGenet shows associations with different diseases for both TSC1 and TSC2.

**Fig 6:**
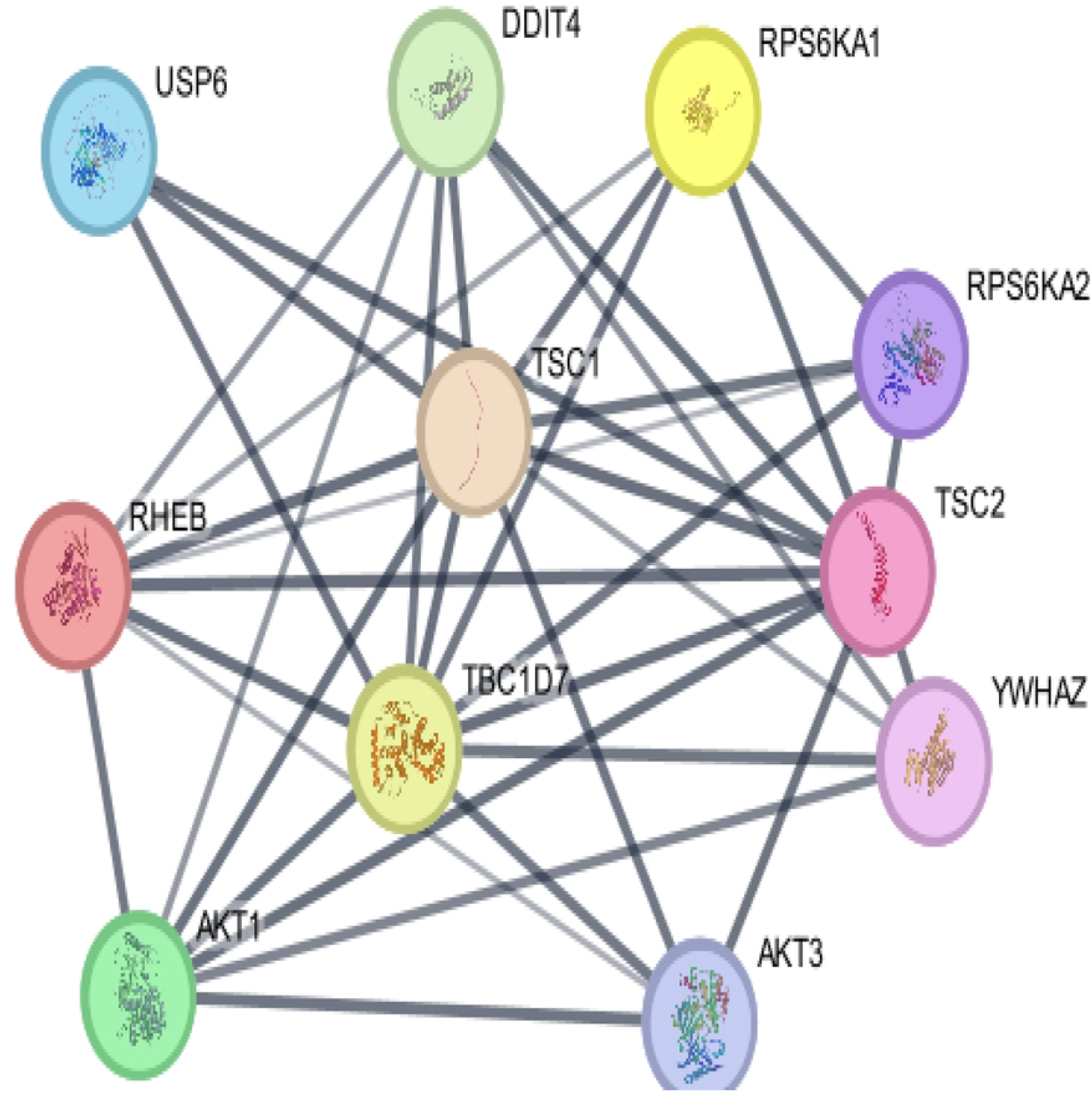
STRING analysis. STRING server shows both TSC1 and TSC2 with RHEB, DDIT4, YWHAZ, RPS6KA1, RPS6KA2, AKT3, AKT1, USP6, and TBC1D7.

Both proteins exhibit high connectivity within this network, interacting with ten other proteins to form a highly interconnected network with 37 edges and 11 nodes. The mean node degree stands at 6.73, while the PPI enrichment value is 4.98e-08. This network suggests these proteins have strong functional associations and potential regulatory relationships.

### 3.10 Superimposition of wild-type protein with mutant structure

Structural comparison between wild-type and SNP-mutated TSC1 protein using ChimeraX Matchmaker revealed 30% of residues exhibited changes in secondary structure assignment (SS fraction = 0.3), suggesting local alterations in helices, sheets, or coil formations due to SNP-induced amino acid substitutions. Superimposition of TSC1 with SNP missense variant is present in **Fig 7** and of *TSC2* in **Fig 8**.

**Fig 7:**
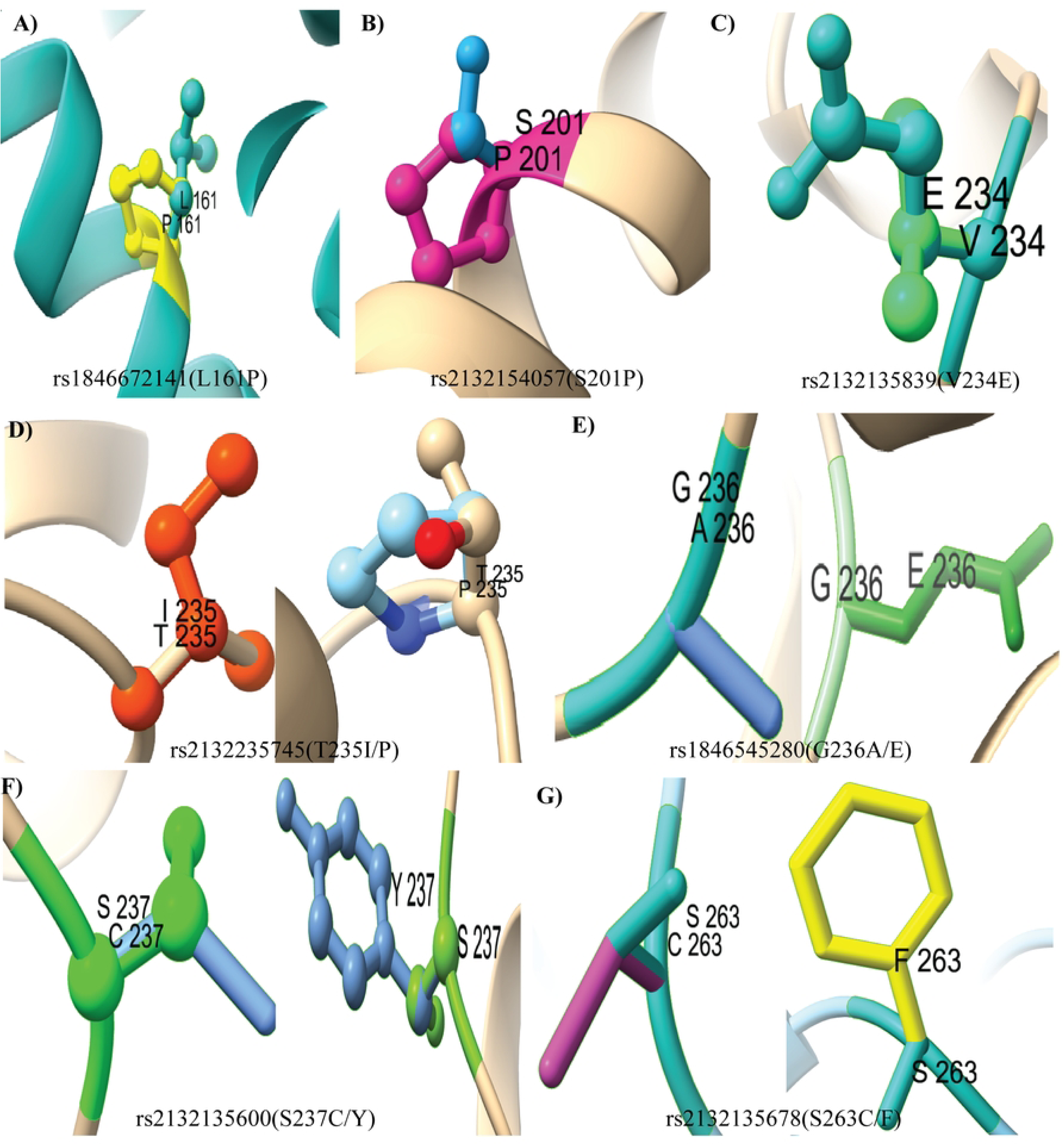
3D model structure of *TSC1* and its nsSNPs. Superimposition of wild-type *TSC1* with variant (A) L161P, (B) S201P, (C) V234E, (D)T235P, I, (E) G236E, A, (F)S237C, Y, (G) S263C, F.

**Fig 8:**
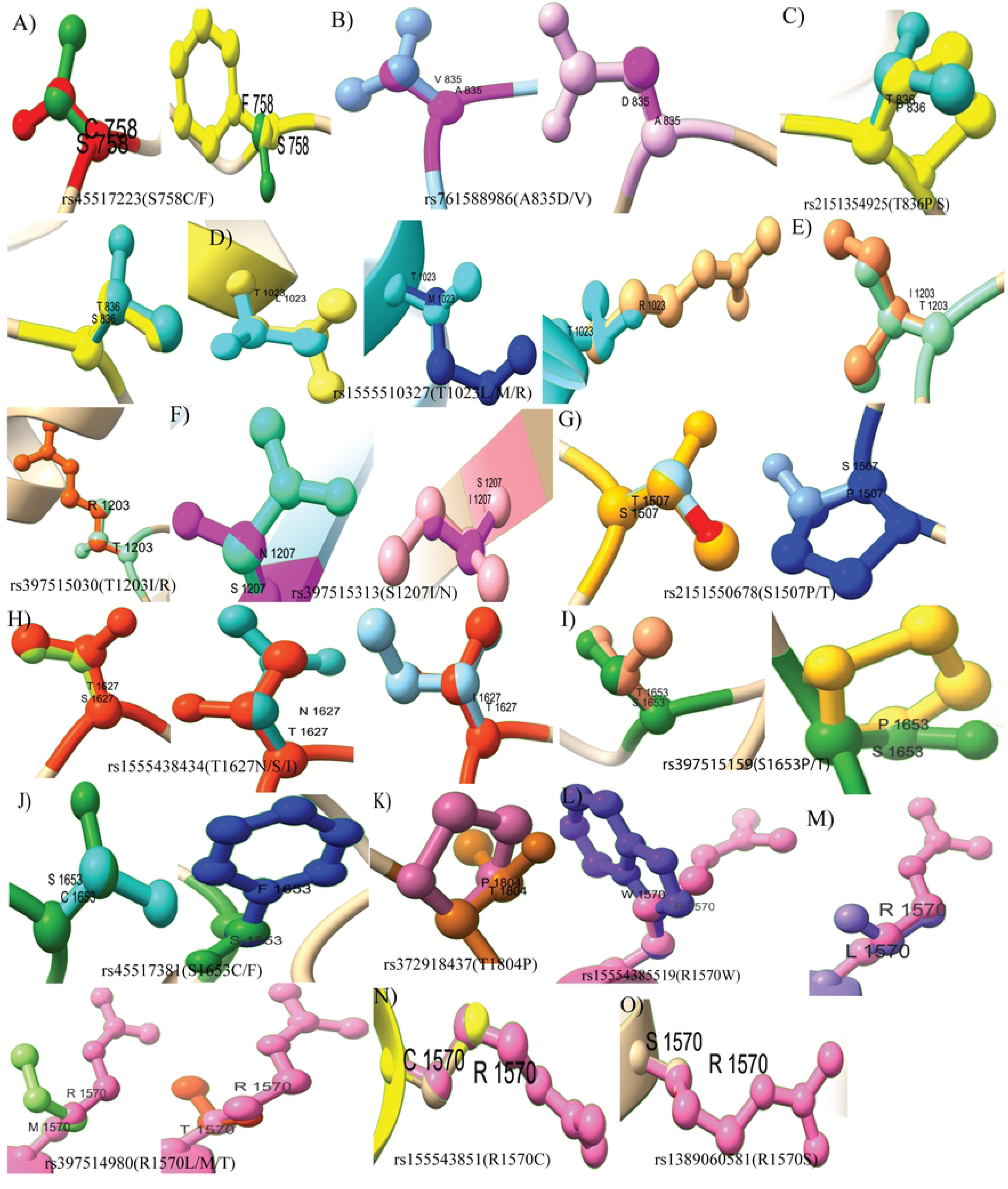
3D model structure of *TSC2* and its nsSNPs. Superimposition of wild-type TSC2 with variant (A) S758C,F, (B) A835V,D, (C) T836P,S, (D)T1023L, M, R, (E) T1203 I, R, (F) S1207I, N, (G) S1507P,T (H) T1627I, N, S, (I) S1653P, T, (J) S1653C, F, (K) T1804P, (L) R1570W (M) R1570L, T, M, (N) R1570C, (O) R1570S.

### 3.11 Molecular Dynamics Simulation

The RMSD value evaluates the average distance between the alpha-carbon backbones of mutant and wild-type models, and a higher RMSD value is associated with a greater deviation of the mutant structure from the wild-type. TSC1 maintains an average RMSD of approximately 0.3-0.35 nm, showing moderate fluctuations. The V234E variant generally shows slightly higher fluctuations compared to the wild type, oscillating around 0.35-0.4 nm, and at times reaching up to 0.45 nm. The S201P variant and L161P variant exhibit similar behavior to the wild type, generally maintaining RMSD values between 0.3 and 0.4 nm, though S201P reaches 0.45 at times. The T235I variant shows slightly higher RMSD values compared to the wild-type and T235P variant. It generally fluctuates between 0.35 and 0.5 nm, indicating a potentially more dynamic or slightly less compact structure. The T235P variant demonstrates RMSD values very close to those of the wild-type TSC1, consistently fluctuating around 0.3-0.35 nm, suggesting comparable structural stability. G236A variant shows very similar RMSD behavior to the wild type, largely overlapping the wild type. G236E variant, however, stands out with noticeably higher RMSD values between 0.35 nm and 0.5 nm, often reaching peaks close to 0.5 nm. S237C variant and S237Y variant both exhibit RMSD fluctuation between 0.3 and 0.4 nm after equilibration, that are very similar to each other and largely consistent with the wild-type protein. S263C and S263F both variant generally fluctuate around 0.35-0.4 nm, showing an upward trend towards the end of the simulation, occasionally reaching above 0.45 nm. **Fig 9(A-E)**. RMSF plot shows the area of greater flexibility due to mutation resulting in disruptive protein folding. L161P, S201P and V234E almost matches wild type TSC1. T235I deviates more than the T235P variant around 225-275 residue. Similarly, the G236E variant also shows more fluctuation in the same area than the G236A variant. Though S237C/Y variants show slight fluctuation around 200-250 residues, S263C/F show almost no deviation from the wild type **Fig 10(A-E)**.

**Fig 9:**
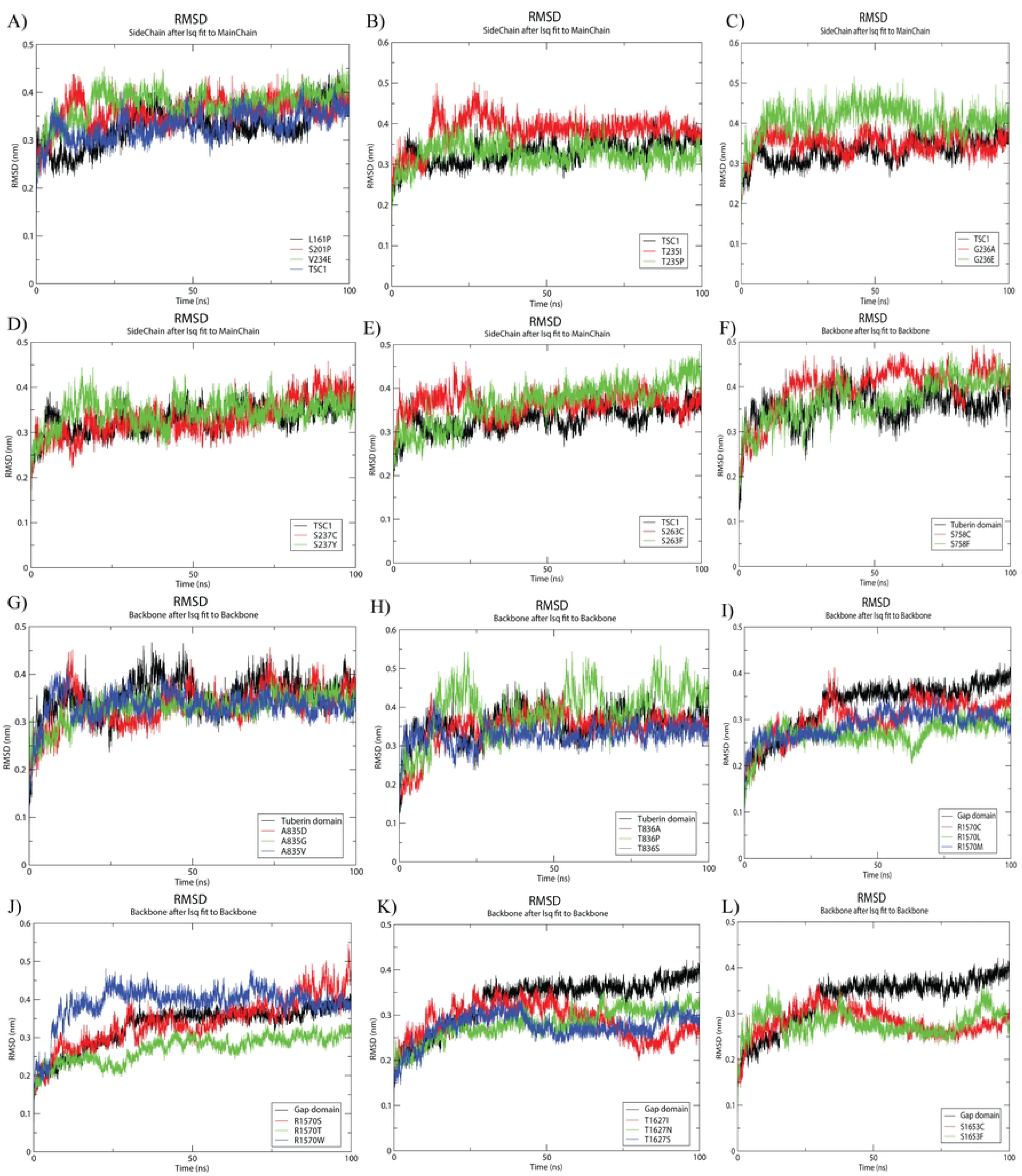
Structural stability analysis of wild-type and SNP variants of TSC1 and TSC2 domains. (A-E) RMSD plot of wild-type Rho-GTPase domain of *TSC1* and SNP. (F-H) RMSD plot of wild-type Tuberin domain of *TSC2* and SNP. (I-L) RMSD plot of the wild-type Gap domain of *TSC2* and SNP.

**Fig 10:**
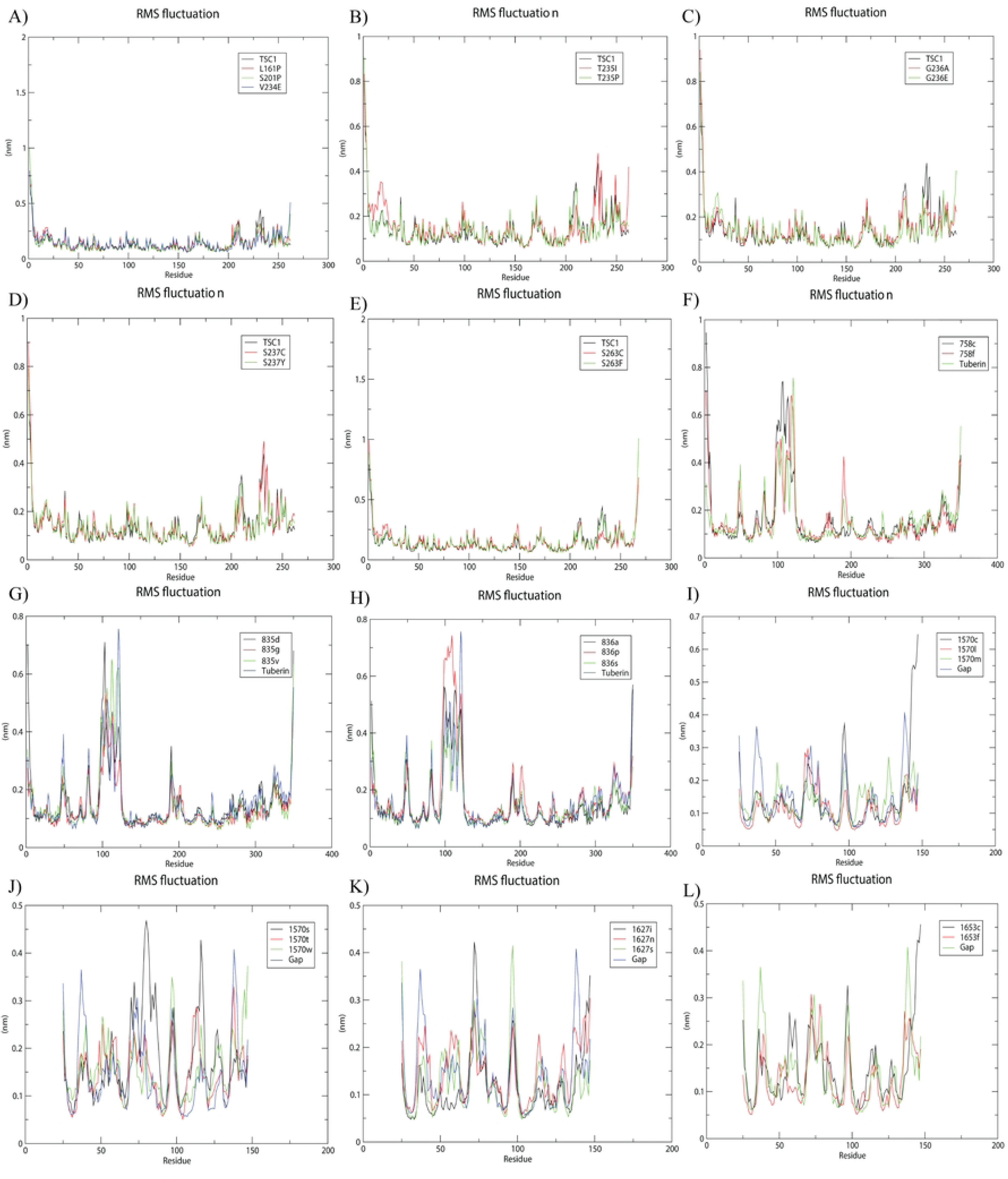
Residue-wise flexibility analysis of wild-type and SNP variants of TSC1 and TSC2 domains. (A-E) RMSF plot of the wild-type Rho-GTPase domain of *TSC1* and SNP. (F-H) RMSF plot of wild-type Tuberin domain of *TSC2* and SNP. (I-L) RMSF plot of the wild-type Gap domain of *TSC2* and SNP.

RMSD of the Tuberin domain is between 0.3 to 0.45. S758C variant fluctuates between 0.4 and 0.5, pointing to structural instability, but the 758F value stays between 0.3 to 0.4. The T836P variant shows a drastic RMSD value from 0.4 to 0.55, but T836A and T836S closely match wild type tuberin domain. A835G, D, and V variants remain between 0.3 to 0.4 **Fig 9(F-H)**. RMSF of the S758C variant shows more flexibility in 100-150 residues but S758F causes flexibility in 200-225 residues. Similarly, T836P varies the most in 100-110 residues though T836A and T836S variants almost overlap wild type tuberin domain. Also, their RMSF plot mostly matches the wild-type tuberin domain **Fig 10(F-H)**. GAP domain shows RMSD value between 0.3 to 0.4. In 1570 position, amino acid R was replaced by C, L, M, S, T, W but only R1570W and R1570S demonstrates RMSD value of 0.4 to 0.5. The others R1570C, R1570L, R1570M, R1570T remain within 0.2 to 0.35 which is less than the wild type. Both T1627I, N, S and S1653F, C variants demonstrate RMSD value of 0.2 to 0.35 **Fig 9(I-L)**. RMSF of both R1570C and R1570W deviates from Gap domain in 100 and 150 residues though R1570S shows deviation in 75 and 120 residues. Also, R1570M deviates in 50 and 100-150 residues. S1653C fluctuates around 60 and 140 residues but S1653F doesn’t deviate much from Gap domain. Though T1627I varies from Gap domain at 75 residues but T1627S shows at 100 residues and T1627N almost matches **Fig 10(I-L)**.

## Discussion

The TSC1 and TSC2 genes have been extensively studied for their role in regulating cell growth, proliferation, and differentiation across various human tissues (53–55). Germline mutations in TSC1 and TSC2 are known to disrupt the formation of the TSC protein complex, which negatively regulates mTORC1 signaling, ultimately contributing to tuberous sclerosis complex (TSC), a multisystem genetic disorder characterized by tumor formation in multiple organs (56,57). However, the in silico understanding of the structural and functional consequences of deleterious non-synonymous single-nucleotide polymorphisms (nsSNPs) in these genes has remained largely unexplored. This project aimed to predict the impact of deleterious nsSNPs in TSC1 and TSC2 at molecular, functional, and structural levels using comprehensive computational approaches to identify the most damaging variants and their potential consequences on the stability and interaction of these proteins.

Twelve different tools were selected to predict most deleterious nsSNPs after evaluating their efficacy via ROC curve (58). Two computational approaches, MuPro and I-Mutant 2.0, were employed to assess the impact of identified nsSNPs on protein stability. Some studies have reported that decreased protein stability induces misfolding, aggregation, and degradation of the protein (59,60) nsSNPs, which were consistently forecasted by both tools to destabilize the protein, were kept for further studies. Highly conserved regions hold critical roles in maintaining the molecule’s function and structure, where changes are often deleterious, while more variable regions tolerate mutations without significantly compromising the macromolecule’s performance (52). Evolutionary conservation analysis revealed that all deleterious nsSNPs are in conserved regions. Twelve “high risk” nsSNPs are found in TSC1 and sixteen in TSC2.

Phosphorylation is a critical regulatory mechanism governing the activity, stability, and subcellular localization of the TSC1–TSC2 complex. TSC2 phosphorylation by AKT promotes mTORC1 activation whereas TSC2 phosphorylation by AMPK inhibits mTORC1 so loss or gain of phosphorylation sites can therefore flip the regulatory outcome (61). NetPhos 3.1 program predicted all deleterious nsSNPs of both TSC1 and TSC2 at phosphorylation sites. In this research, the MutPred2 web server was employed to investigate potential molecular alterations in the structure or function of both TSC1 and TSC2 caused by mutations. According to their g and p scores, every detected deleterious SNP was categorized as “pathogenic. L161P followed by V234E, L90P, G236E, and L264Q showed the highest mutpred score, indicating its significant potential to affect protein structure and function. All SNPs found in TSC1 have also been hypothesized by Mutpred to alter stability and metal binding. As divalent metal ions are essential for proper GAP-domain–mediated GTP hydrolysis (62), alterations affecting metal-binding capacity may directly impair regulation of Rheb-mTORC1 pathway. TSC1 missense substitutions—particularly within the N-terminal/stability region—can reduce hamartin steady-state levels and impair TSC1–TSC2–dependent inhibition of mTORC1 (63,64). In TSC2, S1653P followed by S1207I, S1653F and T1804P had the highest mutpred score. Missense changes in TSC2 (GAP domain) can be functionally deleterious (65). A study annotates S1653 substitutions as key conformational elements of the GAP domain (66). Project HOPE provides new insights into the adverse effects of point mutations on protein structure. It demonstrates notable variances in physicochemical characteristics between wild and mutant amino acids, encompassing size, charge, and hydrophobicity(67). STRING analysis shows that both TSC1 and TSC2 protein plays crucial roles in several vital processes. Their interactions with RHEB, AKT1/3, RPS6KA1/2, and YWHAZ highlight direct control of mTORC1 activity through GAP function (68,69). Association with DDIT4 underscores the role of the TSC complex in stress-induced mTOR suppression (70), while TBC1D7 stabilizes complex assembly (71). Aside from the role of Tuberin in the benign tumors of TSC patients, Tuberin mutations have also been found to cooperate with oncogenic mutations aiding in the initiation and progression of several malignant cancers affecting the brain (medulloblastoma), lung, kidney (renal cell carcinomas), and breast (72–74). Cardiac hypertrophy and type 2 diabetes have also been linked to TSC via AMPK and p70S6K hyperactivation because of Tuberin malfunction(75). As both TSC1 and TSC2 don’t have a crystal structure and are very large in size, we modelled specific domains such as Rho-GTPase domain of TSC1, tuberin and GAP domain of TSC2. nsSNPs such as G236E, S263C/F and V234E with highest fluctuation indicated by both RMSD and RMSF plot, occur in the hamartin region involved in regulating cell adhesion and motility by interacting with the Rho family of GTPases. These could cause loss of adhesion to the extracellular matrix, which correlates with tumor progression in various types of human tumors and is a hallmark of transformation initiated by oncogene activation (76). In TSC2, S758F and T836P showing highest fluctuation present in the tuberin region has N-terminal region, which is crucial for binding to TSC1 (77). MD simulation of the TSC2 GAP domain revealed that R1570W/S exhibit elevated RMSD values, suggesting increased conformational deviation, other substitutions at the same position and variants at T1627 and S1653 displayed RMSD values comparable to or lower than the wild type, reflecting possible increased local rigidity rather than structural preservation (78). RMSF analysis further demonstrated mutation-specific redistribution of flexibility across the domain, indicating long-range allosteric effects (79). These findings suggest that pathogenic TSC2 variants may impair GAP function by altering dynamic properties rather than causing complete structural unfolding (66,80). Till now, missense mutation reported in GAP domains are L1594M, N1643K, N1651S, P1675L, N1681K of TSC patients. Though the SNPs found in this project don’t match the previous evidence, they may reveal new functions on GAP domain. There are also limitations to this study. Computational tools often lack experimental validation, may disagree in their predictions, and are constrained by incomplete structural data for full-length proteins like TSC1, TSC2. Also, silico results cannot account for tissue-specific expression, compensatory mechanisms, or interactions with other regulatory proteins and pathways. Further functional characterization, alongside a complementary clinical study, would be essential to correlate these SNPs with potential changes in protein function or disease phenotypes.

## Acknowledgments

The authors thank the Advanced Bioinformatics Lab, Department of Biochemistry and Molecular Biology, SUST, for providing computational resources and support.

## Author Contributions

Conceptualization, Data Curation, Formal Analysis, Methodology, Software, Visualization, Writing – Original Draft Preparation – TA

Conceptualization, Data Curation, Investigation, Methodology, Project Administration, Supervision, Validation, Writing – Review & Editing - SA

